# Detection of sexually antagonistic transmission distortions in trio datasets

**DOI:** 10.1101/2020.09.11.293191

**Authors:** Elise A. Lucotte, Clara Albiñana, Romain Laurent, Claude Bhérer, Genome of the Netherland Consortium, Thomas Bataillon, Bruno Toupance

## Abstract

Sex dimorphisms are widespread in animals and plants, for morphological as well as physiological traits. Understanding the genetic basis of sex dimorphism and its evolution is crucial for understanding biological differences between the sexes. Genetic variants with sex-antagonistic effects on fitness are expected to segregate in populations at the early phases of sexual dimorphism emergence. Detecting such variants is notoriously difficult, and the few genome-scan methods employed so far have limited power and little specificity. Here, we propose a new framework to detect a signature of sexually antagonistic selection. We rely on trio datasets where sex-biased transmission distortions can be directly tracked from parents to offspring, and allows identifying signal of sexually antagonistic transmission distortions in genomic regions. We report the genomic location and recombination pattern surrounding 66 regions detected as potentially under sexually antagonist selection. We find an enrichment of genes associated with embryonic development within these regions. Last, we highlight two candidates regions for sexually antagonistic selection in humans.

## INTRODUCTION

Males and females primarily differ by the size and number of their gametes, and this asymmetry generates fundamental differences in how fitness is gained in each sex (Parker *et al*. 1972). As a result, a sexual conflict, *i.e*. when selection on a trait acts in opposite direction between the sexes may arise. Genetic variants conferring modifications of a phenotypic trait may be favored in females but disfavored in males and *vice-versa*. These traits are said to be under Sexually Antagonistic (SA) selection. SA genetic variations encoded by the same sets of genes in both sexes lead to an Intralocus Sexual Conflict (IASC). The resolution of IASCs, notably via the evolution of sex-biased expression, is believed to be the primary mechanism for the emergence of sexual dimorphism (Parsch and Ellegren 2013).

Although IASCs have been extensively studied, both theoretically and empirically, many fundamental questions remain unanswered (Mank 2017). In particular, the genomic architecture of these conflicts, *i.e*. their genomic signature, localization and effect on genetic diversity, is subject to debate. Theory predicts that unresolved IASCs influencing survival can lead to a stable polymorphism at SA loci (Rice 1984). It is also expected that female-advantageous alleles are more frequently found in females than in males and *vice-versa* for male-advantageous alleles. However, a substantial difference in allelic frequency between the sexes can only occur if a large number of spontaneous abortion or selective death happen in the population. In humans, while it is unlikely that such selection takes place after birth as mortality during infancy is low in Westernized populations (esa.un.org), strong selection can potentially occur before birth. Indeed, the survival probability of an embryo is estimated to be less than 50% in humans (Benagiano *et al*. 2010) and differences in allelic frequencies between the sexes have been observed among human newborns (Ucisik-Akkaya *et al*. 2010), suggesting that substantial amounts of sex-biased selection may occur before birth.

Previous studies have relied on intersexual F_ST_ to detect ongoing IASC on survival (Cheng and Kirkpatrick 2016; Lucotte *et al*. 2016; Flanagan and Jones 2017; Wright *et al*. 2018), but recent studies argue that this index has limitations. Indeed, high intersexual F_ST_ can be observed in the absence of IASC if selection is limited to one sex or acts with different strengths in each sex. Moreover, it has low power because differences in allelic frequencies between the sexes are expected to be small and a high selection coefficient is needed for them to be detectable (Chippindale *et al*. 2001; Kasimatis *et al*. 2017, 2019).

Theory predicts that the maintenance of polymorphism at a SA locus is facilitated if linked to a distorter locus (Úbeda and Haig 2005; Patten 2014), which would lead to a transmission distortion (TD,*i.e*. non Mendelian transmission of alleles to offspring) occurring before birth either at gamete production (meiotic drive), after copulation (gametic selection) or at fertilization (cryptic choice of the sperm by the ovule). Hence, haplotypes undergoing TD are expected to be enriched for loci with sex-specific effects (Burt and Trivers 2006). Therefore, a locus undergoing IASC is likely to be transmitted in a sex-biased way: parents would transmit more often one allele to their sons and another allele to their daughter either *via* selection on survival during embryonic development or *via* sex-biased TD.

In this study, we propose a new approach to detect signature of IASC based on tracking the transmission patterns of alleles from parents to offspring. We rely on trio datasets and focus on sex-biased TD in offspring. Our method models explicitly the strength and direction of TD and whether it acts in a sex-specific manner, allowing to distinguish different types of sex-biased TD: i) sex-antagonistic: one allele is preferentially transmitted to one sex and the other allele is preferentially transmitted to the other sex, ii)sex-differential: the same allele is preferentially transmitted with different intensities to both sexes and iii)sex-limited: one of the sex is under TD.

We first describe our method. Second, we apply it on the Genome of the Netherlands (GoNL) dataset, which comprises 250 human trios sequenced at 13X coverage (The Genome of the Netherlands Consortium 2014), and explore how widespread IASCs acting on survival are in the human genome. Third, we highlight two candidate regions undergoing sex-antagonistic TD.

## MATERIAL AND METHOD

### Dataset and filtering

The Genome of the Netherland dataset (The Genome of the Netherlands Consortium 2014) comprises 250 parents-child trios (98 Sons and 150 daughters) sequenced at a median coverage of 13X.

We first verified the sex labels by looking at the percentage of X chromosome heterozygosity in males (under 2%) and females (over 6%). For two couples of parents, the males were mislabelled females and *vice-versa* (n°78 and 244).

Only bi-allelic SNPs that passed the quality control of GoNL were retained (Boomsma *et al*. 2014; The Genome of the Netherlands Consortium 2014). The pseudo-autosomal regions on the X chromosome were removed (hg19 positions, chrX:60001-2699520 and chrX:154931044-155260560). X-linked SNPs presenting at least one heterozygous male where removed. Because of the trio structure of the dataset, we were able to test for Mendelian errors, and therefore marked the genotype as missing data for further analyses. Furthermore, SNPs with 2 or more Mendelian errors were removed. At this stage, the dataset comprised 16,980,626 SNPs genome-wide. We removed SNPs with less than 150 informative trios (*i.e*. when at least one parent is heterozygous), for autosomal loci, and 75 informative trios (*i.e*. when the mother is heterozygous) for X-linked loci. In the final dataset, 1,709,245 autosomal SNPs and 50,204 X-linked SNPs were kept.

We verified that the dataset was not genetically structured by sex. We calculated genetic distance matrices in parents for the autosomes and the X chromosome, independently. In this analysis, SNPs in linkage-disequilibrium (r^2^ > 0.25) and individuals with more than 0.5% of missing data were removed. For autosomes, 1 million SNPs were randomly picked 10 times independently and all SNPs were included for the X chromosome. One X chromosome at random was kept for females, and this operation was iterated 30 times independently. Distance matrices between all individuals were calculated using the allele-sharing distance (ASD) and Multi-Dimensional Scaling (MDS) were constructed from those matrices. To determine if male-female distances were significantly different than zero, we performed a Mantel test on the distance matrix between males and females and a matrix where distances between males and females were equal to one and distances between individuals with the same sex were equal to zero. For each repetition, the correlation between both matrices was never significant, either for the autosomes or the X chromosome (Figure S6).

The small difference in age when sampled between males and females should not have an impact on our results for both in parents (median of 61 years in females and 63 years in males) and in children (median of 35 years in females and 34 in males) (Figure S7).

### The likelihood method

We developed a maximum likelihood framework tailored specifically to analyze the transmission of alleles in a set of parents-offspring trios. In our framework, all trios are assumed to be independent (genetically unrelated) and, for a given variant, we only exploit the information brought by informative trios (see Figure 1). We also consider one polymorphic position at a time, although extensions to model the transmission of haplotypes could also be in principle developed.

**Figure 1.**
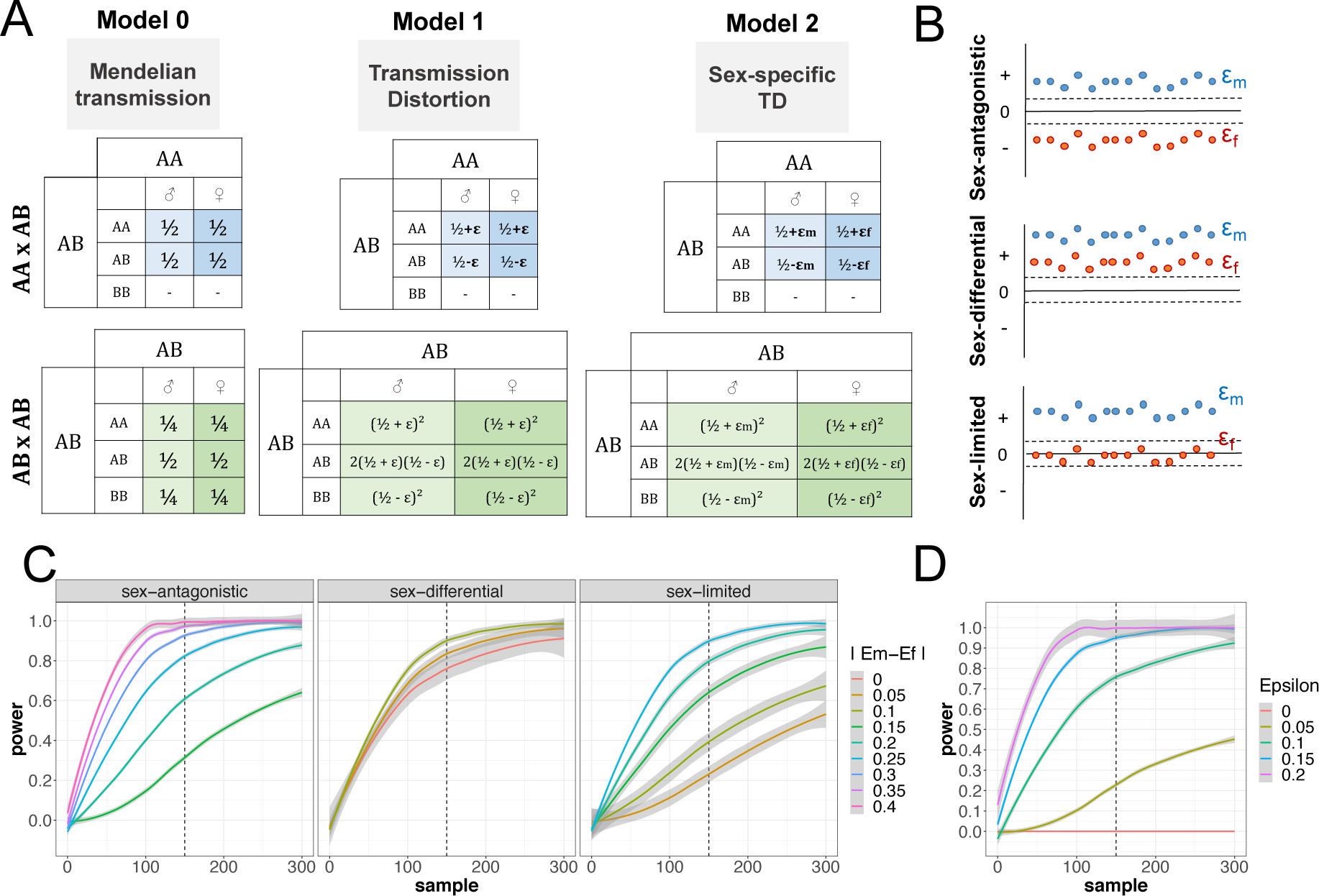
Overview of the likelihood method. **A-** Probability of transmission tables for each model, for AAxAB parents and ABxAB parents. Model 0 is Mendelian transmission, Model 1 is standard transmission distortion, with a unique distortion parameter ε, and Model 2 is sex-specific transmission distortion with sex-specific distortion parameters: ε_m_ for males and ε_f_ for females. **B-** Schematics of the inferred sex-specific distortion parameters in sex-antagonistic, sex-differential or sex-limited regions. **C-** Power simulations at a 0.05 significance level to detect sex-specific TD in case of sex-antagonistic, sex-differential or sex-limited TD, as a function of the number of informative trios and depending on the absolute value of the differences between ε_m_ and ε_f_. **D-** Power simulation at a 0.05 significance level to detect different value of a non-sex-specific ε for different values of ε and number of informative trios.

Within each trio, the transmission of an allele from parents to offspring is modeled using 3 transmission models (*M*_0_, *M*_1_ and *M*_2_ see Figure S1A) that make different assumptions on the effect size and type of transmission distortion affecting a SNP.

At each variable position where at least 150 informative trios are available (75 for the X chromosome), the (natural log) likelihood of the data (ln*L*) under each model is calculated as a series of binomial or multinomial probabilities (see supplementary information for the explicit formulation of the likelihood under each model). The likelihood functions under each transmission model *M*_i_ are maximized analytically thereby yielding (maximum likelihood) estimates for the ε’s as well as measures of statistical uncertainty around the ε estimates (95% approximate confidence interval from likelihood profiles). Last, likelihood ratio tests (LRTs), calculated as differences in deviance between models are used to quantify the amount of statistical support for each alternative model. Note that all three models are nested (*M*_0_, *M*_1_ and *M*_2_) and accordingly the *p*-values associated with each likelihood ratio test statistic was calculated assuming a χ^2^ probability distribution with degrees of freedom calculated as the number of fitted (free) parameters by which the fitted model differ: the LRT between *M*_1_ and *M*_0_ is matched against a χ ^2^ distribution with 1 df, while *M*_2_ versus *M*_0_ is using a χ ^2^ distribution with 2 df.

A local score correction method was used on the p-value of the likelihood of *M*_*2*_ vs *M*_*0*_ to correct for multiple testing (Fariello *et al*. 2017). We used the code made available as a supplementary to this publication. The local scores were computed using the recommended default setting (aggregating p-values p<0.1 yielding a score of 1 or higher).

### Classification into SA, SL and SD TD

Each SNP with a significant p-value of the LRT M2 vs M0 and located in a region enriched in low p-value detected with the Local Score method was classified into SA, SL or SD TD. The decision rule was based on the value of |εm + εf|, for a threshold t of 0.05, which corresponds to the standard deviation of the distribution of epsilons genome-wide.

A SNP is classified as:

- SA if |εm + εf| ≤ maximum (|εm|, |εf|) + t
- SL if |εm + εf|= maximum (|εm|, |εf|) ± t
- SD if |εm + εf| ≥ maximum (|εm|, |εf|) – t

Then, for each regions detected using the Local Score method, if at least 75% of the SNPs could be classified into one category, the region was labelled as this category, otherwise the region was labelled “mixed”.

### Recombination quantile, intersexual F_ST_ and enrichment analysis in candidate regions

To investigate the relationship between presence of sex-specific TD and recombination rate, we downloaded recombination maps from the HapMap phase II project (Frazer *et al*. 2007). We computed the average recombination rate for every autosomal region exhibiting a signal of sex-specific TD. Then, we divided the genome in non-overlapping windows matching the lengths distribution of sex-specific TD regions, and computed the average recombination rate for these windows as a null distribution of genome-wide recombination rates. We used binomial tests to ascertain whether the distribution of recombination rates in sex-specific TD regions matched the null genomic distribution.

For each region type (SA, SL and SD), the intersexual F_ST_ was calculated SNP-wise using the Weir and Cockerham estimator (Weir and Cockerham 1984) and a genome-wide distribution of F_ST_ was computed. Then, we produced empirical null distributions for intersexual F_ST_ by matching an equal number of random genomic regions with comparable heterozygosity and number of SNPs.

We used the refseq genes coordinates from built hg19 to determine which genes were located in the candidate regions. EnrichR was used to perform the functional enrichment analysis, and the tissue-expression enrichment analysis (Chen *et al*. 2013; Kuleshov *et al*. 2016).

### Pipeline for analysing the candidate regions

First, we phased the region using shapeit2 (O’Connell *et al*. 2014). A genetic distance matrix was calculated between all individuals, and a MDS was constructed. Haplotype were identified using a density-based clustering algorithm (package FPC, function dbscan, (Hennig 2019)). Then, we determined the detailed haplotype transmission pattern and assessed significance for sex-specific TD using a Binomial test where H0 is that the probability of transmission does not depend on offspring sex. Third, we analysed the sequence divergence between haplotypes.

To investigate the recombination landscape in TD regions, we used published sex-specific genetic maps (Bherer *et al*. 2017). These maps were built using recombination data from 6 main sources. In total, the combined recombination dataset comprised over 3 million recombination events inferred using genome-wide genotyping data in families and pertaining to over 100,000 meioses. Due to sample ascertainment in the original studies, the female, male and sex-averaged recombination maps are mainly representative of Europeans.

## RESULTS

### A framework for detecting sex specific transmission distortions

We developed a likelihood-based framework to detect sex-biased TD in offspring using trio sequencing (or genotyping) datasets. This method is applied throughout the genome at informative biallelic SNP, *i.e*. SNP with at least one heterozygous parent, examines the fit of the data to three alternative models for the transmission of SNPs from parent to offspring (Figure 1A). All models incorporate a distorsion parameter (ε) that measures the strength and direction of the transmission distortion acting on the alternative allele at a given SNP: ε is zero under Mendelian transmission (model *M*_0_), different from zero but identical in both sexes under classical TD (model *M*_1_), and ε is expected to have sex-specific values (ε_m_ for male offspring and ε_f_ for female offspring) in case of sex-specific TD (Model *M*_2_). A likelihood ratio test (LRT) is performed between *M*_*0*_ and *M*_*2*_ to detect specifically loci under sex-biased TD.

The sex-specific ε parameter estimates are then used to classify a SNPs exhibiting a sex-biased TD signal into sex-antagonistic (SA, *i.e*. TD in both sexes and in opposite direction), sex-limited (SL, *i.e*. TD in only one sex) and sex-differential (SD, *i.e*. TD in both sexes and in the same direction but with different strength) (see methods and Figure 1B). Genomic regions with 75% or more SNPs with one type of TD signal were categorized as such, and those that could not be classified were labelled “mixed” regions.

By modeling explicitly sex-biased TD, we rely on a more specific signal than mere intersexual F_ST_ to track IASCs because we only consider the sub-sample of informative trios (where at least one parent is heterozygote) to evaluate the direction of transmission. Moreover, this method allows to distinguish between SA, SL and SD TD confidently, which is impossible with intersexual F_ST_ because SL and SD selection also leads to higher intersexual F_ST_, nor with the classical Transmission Disequilibrium Test to discover TD (Spielman *et al*. 1993).

We first performed power analyses on simulated trio data to evaluate the ability of our method to detect SNPs that undergo sex-biased TD (Figure 1C-D). We performed two types of simulations, one with equal transmission to male and female offspring and one with sex-specific distorsion parameters, each types with 500 repetitions. We varied the number of informative trios available, the difference between ε_m_ and ε_f_ from 0 to 0.4 (| ε_m_ - ε_f_ |, Figure 1C) and the magnitude of the ε affecting a SNP from 0 to 0.2 (Figure 1D). The power corresponds to the proportion of significant p-values accross repetitions (alpha=5%). As expected, the power increases with the sample size and the effect size. For sex-antagonistic TD, we show that for a sample of 150 informative trios, which is our cutoff, the power is 0.6 for |ε_m_ - ε_f_|= 0.2, 0.8 for |ε_m_ - ε_f_|= 0.25 and 0.95 for |ε_m_ - ε_f_|=0.3 (Figure 1C). Moreover, for 150 informative trios, we have a power of 0.75 to detect an epsilon of 0.1, of 0.9 for an epsilon of 0.15 and 1 for an epsilon of 0.2 (Figure 1D). The cutoff chosen in this study of 150 informative trios provides sufficient power to detect sex-biased TD within the sample size of the GoNL trio dataset. Therefore, the power to detect SA is strongly influenced by the sample size of the trio dataset (Figure 1).

The LRT(*M*_*0*_-*M*_*2*_) p-values we obtained for individual SNPs can be analyzed further using method controlling false discovery rate or a local score method (Fariello *et al*. 2017). By doing the latter, we focus on regions enriched in low p-values, mitigating the common issue of the ‘winner’s curse’ in genome-scan approaches and accounting for the fact that several loci that are physically close to a target of TD share the same signal due to linkage disequilibrium.

Our method can be used for both NGS and array-based genotyping datasets, keeping in mind that regions poorly represented in a SNP genotyping chip will be less likely to be considered significant by the local score method. Below we illustrate our method by applying it to the sequencing GoNL trios dataset (The Genome of the Netherlands Consortium 2014).

### Genomic distribution of sex-biased TD

We analyzed 248 trios from GoNL sequenced at a median coverage of 13X. After filtering for informative trios, we screened a total of 1,709,245 SNPs on autosomes and 50,204 X-linked SNP. Instead of doing a correction for multiple testing, we applied the Local Score method to detect region that are enriched in low p-values. We used a threshold of xi=1, which considers p-values lower than 0.1. We detected 66 SA candidate regions in the GoNL data, including 32 containing genes. Moreover, we detected 168 SL regions, 68 SD regions and 230 mixed regions (Figure 2A, Table 1, Figure S1, Table S1).

**Table 1.** Summary table describing the sex-of-offspring specific TD regions detected

|  |  |  |  | Number of SNPs |  |  |
| --- | --- | --- | --- | --- | --- | --- |
| Type | N region | With genes | Median Length | Median | Min | Max |
| Sex Antagonist | 66 | 32 | 72005 | 66 | 14 | 374 |
| Sex Limited | 168 | 70 | 68578.5 | 77 | 13 | 720 |
| Sex Differential | 68 | 33 | 74368 | 60.5 | 18 | 238 |
| Mixed | 230 | 121 | 98567 | 92.5 | 13 | 1111 |

**Figure 2.**
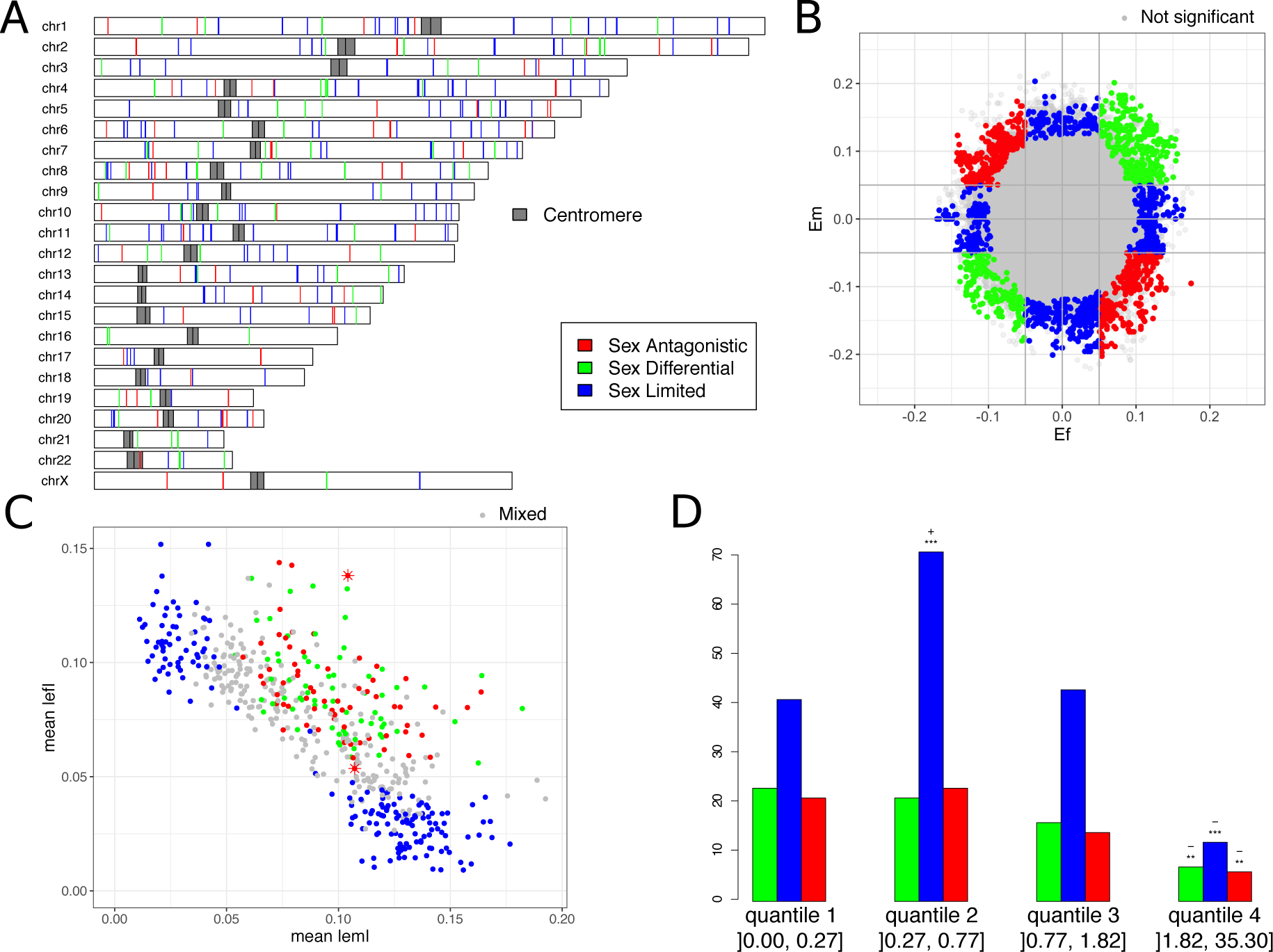
Description of the Sex Antagonistic (Red), Sex Differential (green) and Sex Limited (blue) signal. **A-** Genomic localization of the TD regions. The dark grey rectangles represent centromeres. **B-Classification of the epsilons-** example of chromosome 1. Each point represents one SNP, for which an epsilon m (ε_m_) and epsilon f (ε_f_) were estimated. Grey points represent SNP with a non-significant p-value. **C-Classification of the regions** detected using the Local score method. Each point represents a region, for which a mean value of the absolute ε_m_ and ε_f_ were calculated. The two stars highlight the two regions we analyze further. **D-Number of regions per quantile of recombination rate** (cM/Mb). Each bar represents one type of region. Stars represent the level of significance as measured by a binomial test (H0: 25% of the regions are in each quantile of recombination rate, **: p-value <0.1, ***: p-value <0.01).

We examined the robustness of our findings. First, if we use a more stringent cutoff to aggregate local score (*i.e*. xi=2), we find 38 regions: 11 SA, 12 SD, 7 SL and 8 mixed regions (Table S1). As an alternative, we computed an estimate of the proportion of regions that are coming from Model *M*_*0*_ that posits “no transmission bias” (proportion *π*_0_) *versus* the proportion of region exhibiting some form of distortion (1- *π*_0_). To do so, we first conservatively thinned out SNPs, keeping only SNPs that are 500kb apart to minimize correlation among p-values (n=5272). We obtain a very uniform empirical distribution of p-values across these SNPs, as expected if our LRT is well calibrated and most tests in thinned SNPs are anchored in regions coming from *M*_*0*_ (Figure S2). Using the empirical distribution of thinned p-values, a false discovery rate approach (q-value) estimates that, depending on the cut-off used to estimate *π*_0_, *π*_0_ is 0.98-0.99. This is suggesting that 1- *π*_0_ ≈ 1-2% of the SNPs are anchored in regions that harbour a signal of transmission distortion. This corresponds to roughly 50-10 regions departing from *M*_*0*_. Note however that when assuming a strict FDR approach only one SNP yields a signal that is strong enough to have local FDR< 0.01. This illustrates that more trios are needed to get more precision on the *π*_0_ estimate (as more data will generate clearer separation in the distribution of p-values on SNPs coming from either *M*_*0*_ or alternative SA models such as *M*_2_).

The epsilons values for SNPs classified as SA, SD and SL are displayed on Figure 2B for chromosome 1 as an example. Figure 2C shows the mean absolute values of ε_m_ and ε_f_ for the regions detected as enriched in low p-values using the local score method.

Finally, we investigated the distribution of TD regions with respect to recombination. SA, SL and SD regions are significantly under-represented in the high recombination quantile of the genome (Figure 2B SD p-value=4.41⨯ 10^−3^, SL p-value=3.23⨯ 10^−9^, SA p-value= 2.27⨯ 10^−3^), however this could be due to the Local Score method used to detect regions enriched in low p-values. Indeed, a high recombination rate implies a low LD, which in turns leads to less power to detect significant regions using the Local Score method. SL regions are significantly over-represented in region of medium-low recombination (quantile 2,]0.27, 0.77 cM/Mb], p-value=9.56⨯ 10^−7^).

### Intersexual F_ST_ distributions for the three types of regions

For each set of SA, SL or SD regions, we computed the distribution of intersexual F_ST_ in offspring, and compared it to a matched empirical null distribution of intersexual F_ST_. For each type of TD regions, this null distribution was obtained by randomly sampling genomic regions with matching nucleotide diversity, length and number of SNPs (Figure S3). SA, SL and SD regions show high values of intersexual F_ST_, as compared to matched random genomic regions. Among TD regions, the values for SA regions are significantly higher than both SL and SD (Wilcoxon-Mann-Whitney test, p-values< 2⨯10^−16^). Indeed, for SA, SD and SL regions, the means for the intersexual F_ST_ values in offspring are 0.012 (sd=0.011), 0.000 (sd=0.004) and 0.005 (sd=0.009), respectively. This result is consistent with the expectation that intersexual F_ST_ should be high in regions harboring signals of IASC on survival and that high values can also be detected in case of SL and SD selection (Mank 2017; Wright *et al*. 2018).

### Enrichment analysis

We performed a functional enrichment analysis, focusing on the gene ontologies for biological process and a tissue expression enrichment (Human gene Atlas, GTEx and Jensen tissue, see methods) within the list of genes located in SA, SL and SD regions, using EnrichR (Chen *et al*. 2013; Kuleshov *et al*. 2016). Interestingly, genes present in SA regions are enriched in genes associated with embryonic development (Table 2), both functionally (the growth hormone receptor signaling pathway) and for tissue expression (developmental tissues, *e.g*. placenta, umbilical artery, amniotic fluid). The genes contributing most to enrichment signals are the growth hormone genes (GH2, CSH1, CSHL1, CSH2), which are located in a cluster on chromosome 17, that we will henceforth refer to as the GH locus. Additionally, we find that there is an enrichment in genes that are down-regulated in the uterus and up-regulated in the adipose tissue in females. Genes located in SD and SL regions do not show enrichment in sex-specific functions or development (Table S2 and S3). However, genes located in the mixed regions are expressed preferentially in sex-specific tissues or are enriched in functions related to embryonic development: genes down-regulated in the fallopian tubes and genes involved in embryonic lethality between implantation and somite formation (Table S4).

**Table 2.** Summary of the enrichment found for the list of genes present in SA TD regions. For GO Biological Process and Human Gene Atlas, the enrichment is based on biological functions. For both GTEX and Jensen Tissue databases, the enrichment is based on which tissues the genes are expressed in.

| Database | Term | Overlap | P-value | Adjusted P-value | Z-score | Combined Score | Genes |
| --- | --- | --- | --- | --- | --- | --- | --- |
| GO Biological Process | growth hormone receptor signaling pathway (GO:0060396) | 3/21 | 1.03E-04 | 5.65E-02 | -2.15 | 19.77 | GH2;CSH1;CSHL1 |
| Human Gene Atlas | Placenta | 5/405 | 3.32E-02 | 9.79E-01 | -1.66 | 5.65 | GH2;CSH2;CSH1;OLR1;CSHL1 |
| GTEx Tissue Expression Profile down | GTEx-QCQG-1326-SM-48U24_uterus_female_50-59_years | 4/261 | 2.82E-02 | 1.00E+00 | -1.93 | 6.88 | EPHA10;PPP1R1C;SOD2;ADRA1A |
| GTEx Tissue Expression Profile up | GTEx-0XRO-0226-SM-3LK6F_adipose tissue_female_60-69_years | 11/1039 | 5.79E-03 | 1.00E+00 | -1.84 | 9.48 | BIN3;WTAP;SLC6A16;STRADA;NTRK3;TIMM9;TCP1;DDX42;SOD2;SORBS3;CCAR2 |
| Jensen Tissues | Umbilical_artery | 2/8 | 5.27E-04 | 8.04E-02 | -2.83 | 21.39 | CSH2;CSH1 |
|  | Endometrial_gland | 2/10 | 8.42E-04 | 8.04E-02 | -3.55 | 25.15 | CSH2;CSH1 |
|  | Trophoblast_cell_line | 2/11 | 1.03E-03 | 8.04E-02 | -2.99 | 20.61 | CSH2;CSH1 |
|  | Amniotic_fluid | 3/52 | 1.56E-03 | 9.15E-02 | -3.14 | 20.31 | CSH2;CSH1;THY1 |

### Case study of two potential sexually-antagonistic regions

We identified 32 SA TD regions containing genes (Table S1). We chose to present in details only two regions because one is very strongly contributing to the enrichment signal reported above, and the other is located on the X chromosome, an expected hotspot for the accumulation of SA loci (Rice 1984; Lucotte *et al*. 2016).

#### i. Region on chromosome 17 (chr17:61779927-61988014, 208kb)

This region contains part of the GH locus: CSH1, CSH2, GH2 and CSHL1 (GH1 missing), which are responsible for most of the functional and tissue-expression enrichment in genes located in SA regions (Figure 3, Table 2).

**Figure 3.**
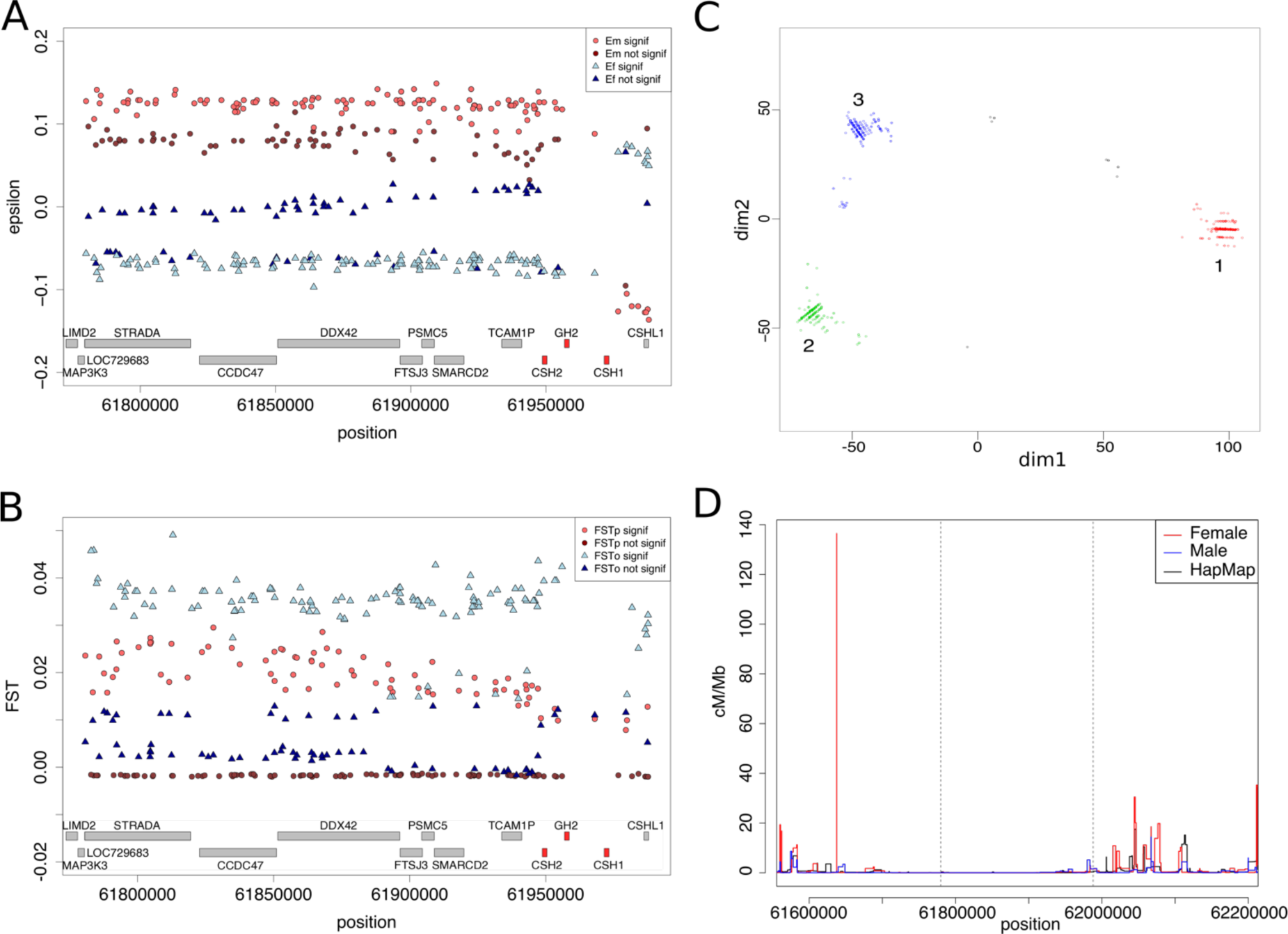
Sexually antagonistic TD region. chr17:61779927-61988014 (208 kb) **A-** Sex-specific distortion coefficients epsilon (ε_m_ in red and ε_f_ in blue). Epsilon differing significantly from zero (LRT p-values lower than 0.05) are in lighter colors. **B-** Intersexual F_ST_ in parents (red) and offspring (blue) for the extended region, significant p-values (as calculated by Fisher exact test, p < 0.05) are in lighter color. **C-** Multi Dimensional Scaling on the genetic distance matrix between all individuals, clustered using a density-based clustering algorithm. Each color denotes a cluster. **D-** The female (red) and male (blue) estimates of recombination rates from our combined genetic maps, as well as the sex-averaged recombination rates (black) from the HapMap linkage genetic map. Horizontal grey lines represent the position of the region of interest.

The absolute values of the εs are comprised between 0.032 and 0.149 (mean 0.107) for males and 0.00 and 0.097 (mean 0.054) for females, with a mean delta| ε_m_ - ε_f_ | of 0.158 (Figure 3A). Note that the inversion of the signs of the epsilons in the vicinity of CSHL1 gene is due to a different attribution of reference and alternative alleles and does not affect our pipeline to detect SA regions. High intersexual F_ST_ values are observed in this 208kb region, both in children and parents (Figure 3B). However, SNPs located next to each other can harbor low and high intersexual F_ST_, suggesting a complex genomic architecture. We phased this region and discovered three distinct haplotypes (Figure 3C and S4). The three haplotypes are almost equally distributed in children (1: 35.8%, 2: 33.8%, 3: 28.2%) and in parents (1: 35.1%, 2: 29.1%, 3: 34.1%). In parents, haplotype 2 is carried by fewer males than expected at random (41.38 % males, p_0_ = 50.20 %, Binomial test p-value = 0.002), while haplotype 3 is carried by an excess of males (57.35 % males, p_0_ = 50.20 %, Binomial test p-value = 0.01). In children, we found an excess of male with haplotype 1 (48.04 % males, p_0_ = 39.20 %, Binomial test p-value = 0.02) while haplotype 2 is still carried by fewer males than females, although not significantly (31.95 % males, p_0_ = 39.20 %, Binomial test p-value= 0.06). Transmissions of these three haplotypes seem to be sex-antagonistic (Table S5): if a parent is heterozygous for haplotype 1 and 3, haplotype 1 is more often transmitted to sons (Fisher exact test, p-value=3.5 7 ⨯ 10^−2^) and haplotype 3 to daughters (Fisher exact test, p-value=4.79 ⨯ 10^−2^). Additionally, if the heterozygous parent has haplotype 1 and haplotype 2 or 3, haplotype 1 is more often transmitted to sons (Fisher exact test, p-value = 3.84 ⨯ 10^−3^) and haplotype 2 or 3 to daughters (Fisher exact test, p-value=4.48 ⨯ 10^−2^). The sample sizes are small, and these p-values do not resist correction for multiple testing, except for the biased transmission of haplotype 1 to sons compared to 2 or 3 (p-value = 3.84×10^−2^ after Bonferroni correction). These results suggest that haplotype 1 is beneficial for males, and deleterious for females.

This region encompasses 1291 SNPs, and has a length of 208,087bp. The mean number of differences between genomics regions are 167.66 SNPs for haplotypes 1 and 2, 151.78 SNPs for haplotypes 1 and 3 and 84.59 SNPs for haplotypes 2 and 3 (Table S6). For such a short region, this suggests that recombination is rare between the three haplotypes. Indeed, this region is a cold-spot of recombination (Figure 3D), flanked by two sex-specific hotspots of recombination.

This pattern is not due to a mapping artifact, either sex-specific or region-specific (Supplementary text II). Moreover, while we could not replicate the transmission results in another dataset because we do not have access to a dataset with enough trios, we were able to replicate the finding of the three haplotypes in other European populations (Supplementary text III).

#### ii. Region on the X chromosome (chrX:47753028-47938680, 186kb)

This region contains the gene SPACA5, the only gene from the SPACA family (5 genes) located on the X chromosome (Figure 4). It encodes a sperm acrosome associated protein; which is directly involved in gamete fusion.

**Figure 4.**
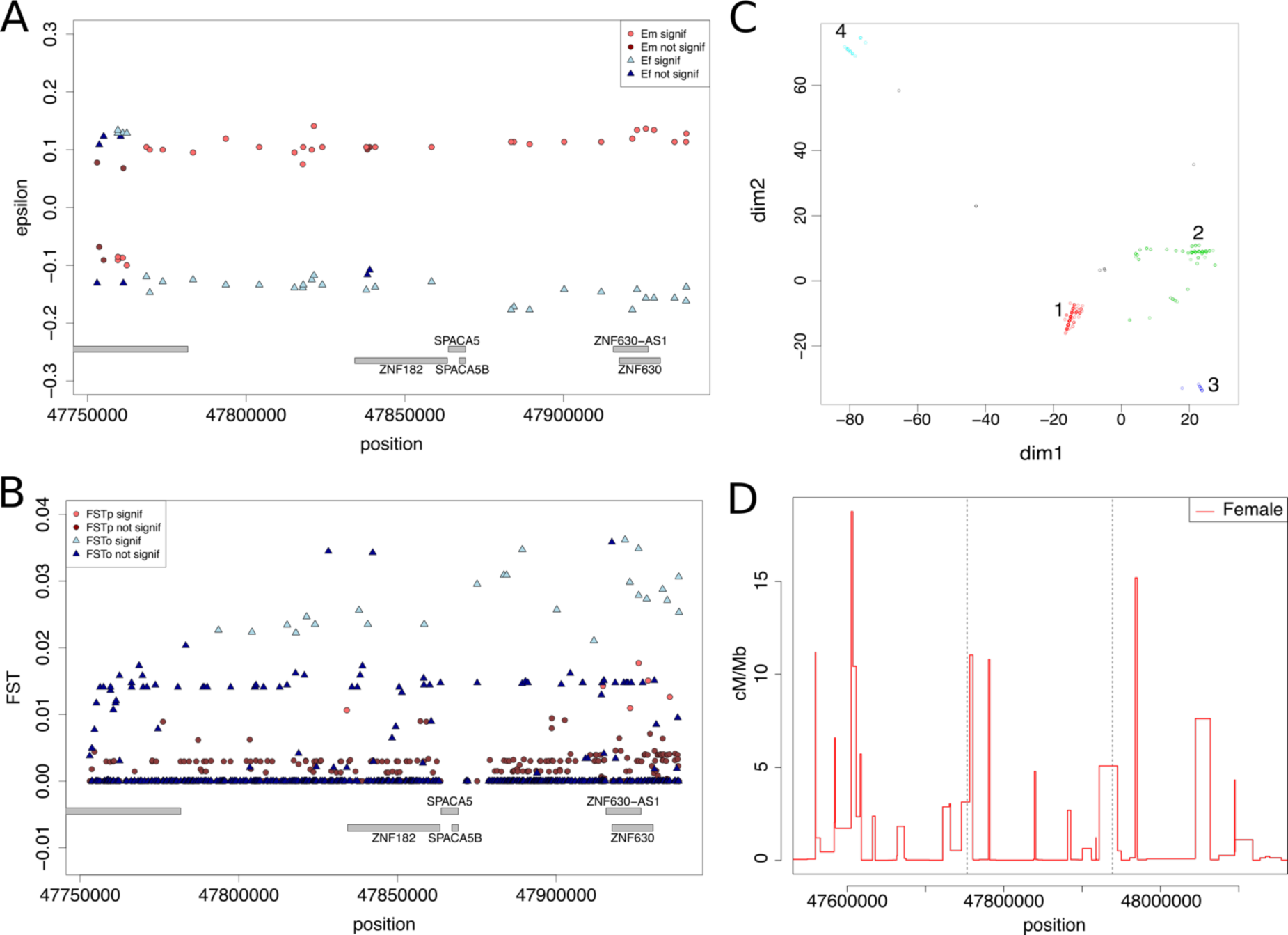
Sexually antagonistic SoO specific TD region. chrX:47753028-47938680 (186 kb) **A-** Sex-specific epsilon (Em in red and Ef in blue). Epsilon with LRT p-values lower than 0.05 are in lighter colors. **B-** intersexual F_ST_ in parents (red) and offspring (blue) for the extended region, significant p-values (as calculated by Fisher exact test, p < 0.05) are in lighter color. **C-** Multi Dimensional Scaling on the genetic distance matrix between all individuals, clustered using a density-based clustering algorithm. **D-** The female (red) estimates of recombination rates from our combined genetic maps. Horizontal grey lines represent the position of the region of interest.

The absolute values of ε’s are comprised between 0.068 and 0.141 (mean = 0.104) for males and 0.108 and 0.177 for females (mean = 0.138) and the mean |ε_m_-ε_f_| is 0.242 (Figure 4A). High intersexual F_ST_ can be observed along the whole region in offspring, and at the right end of the region for parents (Figure 4B). The same pattern of alternating high and low F_ST_ for adjacent SNPs, similar to the region of chromosome 17 highlighted above, was observed. We discovered 4 haplotypes, including two in high frequency: haplotype 1 and haplotype 2 with a frequency of 0.516 and 0.441 in parents (0.528 and 0.428 in children), respectively (Figure 4C, S5). Haplotype 3 and 4 have a frequency of 0.013 and 0.025 in parents (0.012 and 0.026 in children), respectively. Transmission of these haplotypes is significantly sex-biased (Table S7): haplotype 1 is more often transmitted to daughters (Fisher exact test p-value = 2.95×10^−2^) while haplotype 2 is more often transmitted to sons (Fisher exact test p-value = 2.05×10^−2^). Interestingly, when fathers have haplotype 1, mothers are more likely to transmit haplotype 1 to daughters (24 cases against 10 cases of mothers transmitting haplotype 2, Binomial test p-value=2.43×10^−2^). Haplotype 1 and 2 have a lower percentage of divergence in sequence than what we observe for the region on chromosome 17 (Table S8), which can be explained by the occurrence of recombination within the region (Figure 4D).

As above, validation analyses suggest that this pattern is not due to a mapping artifact (Supplementary text II). European populations from the 1000 Genomes project display a similar haplotype structure (Supplementary text III).

## DISCUSSION

We propose a new method to detect sex-of-offspring-specific TD, hereafter referred to as sex-biased TD, using sequencing or genotyping of trio (parent-offspring) datasets to track directly the transmission of alleles in each sex. This offers a way to categorize different types of genomic regions: Sex-Antagonistic (SA), Sex-Limited (SL) and Sex-Differential (SD) TD by providing estimates of the intensity of sex-specific TD. This method circumvents the limitation of previous methods relying solely on intersexual F_ST_, by specifically detecting loci undergoing SA TD. Moreover, by using a Local Score method (Fariello *et al*. 2017), we detect genomic windows with an enrichment in low p-values, as expected under TD, and hence reduce the risk to detect false positives as compared to single locus F_ST_ measurements.

Loci undergoing IASC are expected to experience balancing selection, because different alleles are beneficial in different sexes (Connallon and Clark 2014). It has been proposed to use Tajima’s D, a statistic summarizing the site frequency spectrum (essentially capturing the amount of rare versus frequent alleles), in combination with intersexual F_ST_ to distinguish SA selection from other sex-biased selections (Wright *et al*. 2018). However, signatures of balancing selection are notoriously difficult to detect (Rowe *et al*. 2018). Moreover, in our case, because we only keep SNPs with at least 150 heterozygous trios (75 for the X chromosome), we have an ascertainment bias towards SNPs with an elevated Tajima’s D, whether they show a signal of TD or not.

The power of the method to detect regions with distortions is strongly dependent on the number of informative trios. When analyzing the GoNL trios we only have statistical have power to test SNPs with intermediate frequencies. We expect SNP under SA selection to be at intermediate frequencies (Mank 2017) and to exhibit a large difference in sex-specific distortion parameter, as measured by (|ε_f_-ε_m_|), so SNPs with the strongest amount of SA TD are specifically captured by our method. Although GoNL is one of the largest trio datasets published to date, the limited number of trios precludes from over-speculating on specific regions. In this study, we draw conclusions on overall patterns of SA TD, and merely focus on two regions that seemed the most striking and interesting examples of SA TD.

We detected 66 SA TD regions genome-wide, including 32 with genes. We found that regions undergoing SA TD are enriched for SNPs with high intersexual F_ST_, which is expected (Lucotte *et al*. 2016; Mank 2017). Regions undergoing SL and SD TD also show high intersexual F_ST_, but have significantly lower intersexual F_ST_ than SA TD regions.

We performed a functional and tissue-expression enrichment analysis on the genes located within the SA region. The enrichment analyses performed in SA regions reveal that these contain genes that are primarily involved in developmental functions, and expressed in tissues involved in development. The functional enrichment and expression enrichment were not significant after correction for multiple testing. However, this result is in concordance with the expectation that SA TD may occur during gamete fusion and embryo development.

We then focused on two SA TD regions: a region on chromosome 17 containing the genes responsible for most of the genome-wide enrichment in developmental tissues and the unique SA region containing genes detected on the X chromosome. In both regions, we detected several haplotypes that are preferentially transmitted to one sex or the other, which is in concordance with the prediction of theoretical works (Úbeda and Haig 2005; Burt and Trivers 2006; Patten *et al*. 2010; Ubeda *et al*. 2011; Patten 2014). The chromosome 17 region encompasses the growth hormone locus, notably the GH gene, which encodes a protein in the placenta that is important for *in utero* development (Oberbauer 2015), and affects adult traits such as height and bone mineral density (Timasheva *et al*. 2013). Interestingly, there is evidence for ongoing IASC on human height (Stulp *et al*. 2012). The high sequence divergence among the three haplotypes is probably due to the lack of recombination in this region. Although sample sizes are low, a pattern of SA TD of the haplotypes can be detected. However, the p-values for sex differences in haplotype transmission are nonsignificant.

The X chromosome region encompasses the only SPACA gene on this chromosome, which is expressed in the spermatozoid acrosome, involved in gamete fusion. This is an interesting feature as TD could happen at gamete fusion. Deeper investigations of the role of this gene and the impact of the observed genetic polymorphism are warranted.

We were able to replicate the finding of the number of haplotypes in European populations of the 1000 Genomes dataset, however, a trio dataset of at least equal sample size should be investigated in the future to validate the TD pattern detected in the GoNL data. In the near future, we expect more datasets with pedigrees (trios or extended sibships), on which this method could be used to gain more knowledge on the architecture of SA TD in the human genome.

TD can be due to several non-exclusive mechanisms: after birth and haploid selection, occurring between gamete formation and fertilization or sexually antagonistic selection on survival occurring between fertilization and birth (during embryonic development). Our method does not allow to distinguish between these biological mechanisms. One perspective of this study would be to modify the method to take into account the sex of the parent in TD, which could allow to distinguish between TD occurring before and after fertilization. Indeed, variation in expression profile of the genes in haploid sperm among a single ejaculate has been shown to correlate with motility and fertility in humans, which is consistent with gametic selection happening in humans (Lambard *et al*. 2004).

### CONCLUSION

We provide a new framework to detect loci specifically undergoing sex-antagonistic TD in genomic datasets. It allows to discriminate between sex-antagonistic, sex-limited and sex-differential TD. This circumvents limitations of the intersexual F_ST_ used in previous studies. We detect 32 gene coding regions undergoing sex-antagonistic TD in a human population from the Netherland and highlight two intriguing candidate regions. Our method can be applied to any sequencing or genotyping datasets structured in parents-offspring trios, and constitute therefore an important progress to elucidate the genomic architecture of intralocus sexual conflicts and their implications in sex dimorphisms evolution. As costs of sequencing and genotyping are rapidly decreasing, we expect pedigrees datasets to become commonplace in the future.

## ACKNOWLEDGEMENTS

Part of this work was supported by funding from the Danish Council for Independent Research and the French Ministry for Higher Education and Research. Most of the computing for this project was performed on the GenomeDK cluster. We would like to thank GenomeDK and Aarhus University for providing computational resources and support that contributed to these research results. This study makes use of data generated by the Genome of the Netherlands Project. A full list of the investigators is available from www.nlgenome.nl. Funding for the GoNL project was provided by the Netherlands Organization for Scientific Research under award number 184021007, dated July 9, 2009 and made available as a Rainbow Project of the Biobanking and Biomolecular Research Infrastructure Netherlands (BBMRI-NL). The sequencing was carried out in collaboration with the Beijing Institute for Genomics (BGI).

## AUTHOR CONTRIBUTIONS

EL, BT and TB conceived and designed the study, and acquired funding. This work was supervised by TB and BT. CA and BT formalized and CA coded the likelihood method designed by TB and EL. EL and RL curated and analysed the data. Additional analyses on recombination were performed by CB. EL drafted the initial version of the manuscript and BT, RL, TB and CA contributed to later versions of the manuscript. The Genome of the Netherland Consortium provided the data.

## DATA ACCESSIBILITY

This project was approved by the GoNL Data Access Committee (application nr 2014053).

