## Supplementary_materials for "Detection of sexually antagonistic transmission distortions in trio datasets"

### SUPPLEMENTARY TEXT

Complete list of authors from Genome of the Netherland Consortium:

Dorret I Boomsma, Cisca Wijmenga, Eline P Slagboom, Morris A Swertz, Lennart C Karssen, Abdel Abdellaoui, Kai Ye, Victor Guryev, Martijn Vermaat, Freerk van Dijk, Laurent C Francioli, Jouke Jan Hottenga, Jeroen F J Laros, Qibin Li, Yingrui Li, Hongzhi Cao, Ruoyan Chen, Yuanping Du, Ning Li, Sujie Cao, Jessica van Setten, Androniki Menelaou, Sara L Pulit, Jayne Y Hehir-Kwa, Marian Beekman, Clara C Elbers, Heorhiy Byelas, Anton J M de Craen, Patrick Deelen, Martijn Dijkstra, Johan T den Dunnen, Peter de Knijff, Jeanine Houwing-Duistermaat, Vyacheslav Koval, Karol Estrada, Albert Hofman, Alexandros Kanterakis, David van Enckevort, Hailiang Mai, Mathijs Kattenberg, Elisabeth M van Leeuwen, Pieter B T Neerincx, Ben Oostra, Fernanodo Rivadeneira, Eka H D Suchiman, Andre G Uitterlinden, Gonneke Willemsen, Bruce H Wolffenbuttel, Jun Wang, Paul I W de Bakker, Gert-Jan van Ommen & Cornelia M van Duijn

**I-** **The likelihood method- Derivation of the mathematical formulas**

**II-** **Is the sex-specific pattern of transmission due to a mis-mapping of the reads?**

1. **BLAST analysis**
2. **Analyses of the coverage**

**III-** **Replication in the 1000Genomes populations**

### Derivation of mathematical formulas

Let us consider, in a diploid species, a biallelic locus with alleles $A$ and $B$. We further assume that the genotypes are available for a set of trios consisting of two parents that are both genotyped and a single offspring (although in principle that scheme can be extended to more general sets of sibships than Parent/Offspring trios). After reproduction, transmitted (T) and non-transmitted (NT) allele counts can be arranged in a T-NT matrix (table 1) following Spielman et al.[1]

| | $A_{\mathrm{NT}}$ | $B_{\mathrm{NT}}$ |
|---|---|---|
| $A_{\mathrm{T}}$ | $a$ | $b$ |
| $B_{\mathrm{T}}$ | $c$ | $d$ |

Table 1: T-NT matrix for a biallelic locus in a diploid species

When considering all possible genotypes for parents and offspring, the T-NT matrices are given in table 2.

| parents | $AA \times AA$ | $AA \times AB$ | | $AA \times BB$ | $AB \times AB$ | | | $AB \times BB$ | | $BB \times BB$ |
|---|---|---|---|---|---|---|---|---|---|---|
| offspring | $AA$ | $AA$ | $AB$ | $AB$ | $AA$ | $AB$ | $BB$ | $AB$ | $BB$ | $BB$ |
| T-NT matrix | $\begin{matrix} 2 & 0 \\ 0 & 0 \end{matrix}$ | $\begin{matrix} 1 & 1 \\ 0 & 0 \end{matrix}$ | $\begin{matrix} 1 & 0 \\ 1 & 0 \end{matrix}$ | $\begin{matrix} 1 & 0 \\ 0 & 1 \end{matrix}$ | $\begin{matrix} 0 & 2 \\ 0 & 0 \end{matrix}$ | $\begin{matrix} 0 & 1 \\ 1 & 0 \end{matrix}$ | $\begin{matrix} 0 & 0 \\ 2 & 0 \end{matrix}$ | $\begin{matrix} 0 & 1 \\ 0 & 1 \end{matrix}$ | $\begin{matrix} 0 & 0 \\ 1 & 1 \end{matrix}$ | $\begin{matrix} 0 & 0 \\ 0 & 2 \end{matrix}$ |
| # occurences | $x_1$ | $x_2$ | $x_3$ | $x_4$ | $x_5$ | $x_6$ | $x_7$ | $x_8$ | $x_9$ | $x_{10}$ |

Table 2: T-NT matrix and counts notations for all possible genotypes for parents and offsprings. The last row corresponds to the number of trios in each configuration observed in the data (a total of $n$ trios).

For a given locus, combining tables 1 and 2 over multiple trios, we obtain:

$$\begin{aligned} a &= 2x_1 + x_2 + x_3 + x_4 \\ b &= x_2 + 2x_5 + x_6 + x_8 \\ c &= x_3 + x_6 + 2x_7 + x_9 \\ d &= x_4 + x_8 + x_9 + 2x_{10} \end{aligned}$$

Moreover, the total number of heterozygous parents $n_{het}$ is given by:

$$\begin{aligned} n_{het} &= x_2 + x_3 + 2x_5 + 2x_6 + 2x_7 + x_8 + x_9 \\ &= b + c \end{aligned}$$

If we note $\mathbb{P}[A]$ and $\mathbb{P}[B] = 1 - \mathbb{P}[A]$ the probability that an heterozygous parent transmits to his offspring allele $A$ and $B$ respectively, we can write the likelihood of the data as:

$$\begin{aligned} L =& \binom{x_2 + x_3}{x_2} \mathbb{P}[A]^{x_2} \ \mathbb{P}[B]^{x_3} \\ \times& \binom{x_8 + x_9}{x_8} \mathbb{P}[A]^{x_8} \ \mathbb{P}[B]^{x_9} \\ \times& \binom{x_5 + x_6 + x_7}{x_5, x_6, x_7} \mathbb{P}[A]^{2x_5} \ (2\mathbb{P}[A]\,\mathbb{P}[B])^{x_6}\,\mathbb{P}[B]^{2x_7} \end{aligned}$$

where $\binom{\cdots}{\cdots}$ and $\binom{\cdots}{\ldots,\ldots,\ldots}$ denote respectively the binomial and multinomial coefficients.

Ignoring all terms unlinked to the parameters $\mathbb{P}[A]$ and $\mathbb{P}[B]$, we obtain:

$$
\begin{aligned}
L &= \mathbb{P}[A]^{x_2+x_8+x_6+2x_5} \ \mathbb{P}[B]^{x_3+x_9+x_6+2x_7} \\
&= \mathbb{P}[A]^b \ \mathbb{P}[B]^c \\
&= p^b(1-p)^{n_{het}-b}
\end{aligned}
$$

which is a binomial form using notation $p = \mathbb{P}[A]$, $1-p = \mathbb{P}[B]$ and noting that $c = n_{het} - b$.

### Model 0: Mendelian transmission

In this case:

$$\mathbb{P}[A] = \mathbb{P}[B] = \frac{1}{2}$$

Therefore:

$$L_0 = \left(\frac{1}{2}\right)^{b+c} = \left(\frac{1}{2}\right)^{n_{het}}$$

$$\ln L_0 = -n_{het}\ln(2)$$

### Model 1: Transmission distorsion

Here, one allele is preferentially transmitted to offsprings, which we parametrized by a deviation $\varepsilon$ from the expected $\frac{1}{2}$ probability under Mendelian transmission, *i.e.*:

$$\mathbb{P}[A] = \frac{1}{2} + \varepsilon \quad \text{and} \quad \mathbb{P}[B] = \frac{1}{2} - \varepsilon \quad \text{with} \quad \varepsilon \in \left[-\frac{1}{2}, \frac{1}{2}\right]$$

Therefore:

$$L_1(\varepsilon) = \left(\frac{1}{2}+\varepsilon\right)^b \left(\frac{1}{2}-\varepsilon\right)^c = \left(\frac{1}{2}\right)^{n_{het}} (1+2\varepsilon)^b\,(1-2\varepsilon)^c$$

$$\ln L_1(\varepsilon) = -n_{het}\ln(2) + b\ln(1+2\varepsilon) + c\ln(1-2\varepsilon)$$

In this case, the maximum likelihood estimator for $\varepsilon$ is:

$$\varepsilon_{\text{MLE}} = \frac{1}{2}\frac{b-c}{b+c} = \frac{b-c}{2n_{het}}$$

and loglikelihood at $\varepsilon_{\text{MLE}}$ is:

$$\ln L_1 = -n_{het}\ln(n_{het}) + b\ln(b) + c\ln(c)$$

### Model 2: Sex of offspring specific transmission distorsion

In this model, we introduce sex specific parameters $\varepsilon_m$ and $\varepsilon_f$ for male and female offspring respectively (we also note that the same formalism could be also used to formulate models of sex of parent specific transmission). Counting separately T-NT matrix by considering the sex of

| | $A_{\text{NT}}$ | $B_{\text{NT}}$ |
|---|---|---|
| $A_{\text{T}}$ | $a_m$ | $b_m$ |
| $B_{\text{T}}$ | $c_m$ | $d_m$ |

(a) For male offspring

| | $A_{\text{NT}}$ | $B_{\text{NT}}$ |
|---|---|---|
| $A_{\text{T}}$ | $a_f$ | $b_f$ |
| $B_{\text{T}}$ | $c_f$ | $d_f$ |

(b) For female offspring

Table 3: T-NT matrices for sex of offspring specific model

offspring in each trio, it is then straightforward to rewrite sex-specific tables (see tables 3a and 3b). Following a derivation similar to above we obtain:

$$\begin{aligned} L_2(\varepsilon_m, \varepsilon_f) &= \left(\frac{1}{2} + \varepsilon_m\right)^{b_m} \left(\frac{1}{2} - \varepsilon_m\right)^{c_m} \left(\frac{1}{2} + \varepsilon_f\right)^{b_f} \left(\frac{1}{2} - \varepsilon_f\right)^{c_f} \\ &= \left(\frac{1}{2}\right)^{n_{het}} (1 + 2\varepsilon_m)^{b_m} (1 - 2\varepsilon_m)^{c_m} (1 + 2\varepsilon_f)^{b_f} (1 - 2\varepsilon_f)^{c_f} \end{aligned}$$

$$\ln L_2(\varepsilon_m, \varepsilon_f) = -n_{het} \ln(2) + b_m \ln(1 + 2\varepsilon_m) + c_m \ln(1 - 2\varepsilon_m) + b_f \ln(1 + 2\varepsilon_f) + c_f \ln(1 - 2\varepsilon_f)$$

Maximum likelihood estimators for $\varepsilon_m$ and $\varepsilon_f$ are:

$$\varepsilon_{m_{\text{MLE}}} = \frac{1}{2} \frac{b_m - c_m}{b_m + c_m}$$

$$\varepsilon_{f_{\text{MLE}}} = \frac{1}{2} \frac{b_f - c_f}{b_f + c_f}$$

and loglikelihood at $(\varepsilon_{m_{\text{MLE}}}, \varepsilon_{f_{\text{MLE}}})$ is:

$$\ln L_2 = -n_{het,m} \ln(n_{het,m}) + b_m \ln(b_m) + c_m \ln(c_m) - n_{het,f} \ln(n_{het,f}) + b_f \ln(b_f) + c_f \ln(c_f)$$

### II- Is the sex-specific pattern of transmission due to a mis-mapping of the reads?

#### 1- BLAST analysis

For both regions included in the case study, we investigated whether peculiar genomic properties could explain the TD signal we detect. More specifically, sequence similarity between a given region and a sex chromosome could result in mapping errors, eventually leading to the detection of a spurious sex-biased TD signal. To avoid this caveat, we performed a blast analysis as follows. We downloaded the GRCh37 reference genome from the Ensembl database [1]. Then, for each region of interest, we extracted the corresponding sequence and divided it in 10,000 random fragments of 100 bp. We blasted these fragments against the human genome and recorded hits outside the region of interest with at least 95% identity into a set of hits. Finally, we computed the number of hits from this set on each chromosome, and used a binomial test to test whether the number of hits on the sex chromosomes were different than expected by chance, taking into account chromosome lengths. Results are given in Table II.1. For both region, the number of hits is significantly lower than expected at random, given the length of the region.

**Table II.1**: Results of the blast analyses

| Query | target | # hits observed | # hits expected | pvalue |
|---|---|---|---|---|
| chr17, 95% identity | X | 424632 | 496525,8 | 4.44E-323 |
| chr17, 95% identity | Y | 34489 | 188779,7 | 2.96E-323 |
| chrX, 95% identity | Y | 66004 | 179674,3 | 2.96E-323 |

#### 2- Analyses of the coverage

<u>Chromosome 17 region chr17:61779927-61988014</u>
We extracted all positions from chromosome 17 with a filter == PASS. For each position, we computed the mean of the coverage for all males and all females separately. Then we calculated the mean over all positions per bin of 1,000 bp. Within the whole chromosome bins, we filtered bins with a mean coverage outside of the minimum and maximum coverage of the region of interest, to avoid outliers due to specific regions. The mean coverage over 1000bp bins for the region of interest is presented on figure II.1A. The mean coverage is very close for males and females (13.90 and 13.50, respectively) for the region of interest and the whole chromosome (13.93 and 13.55, respectively, figure II.1B).
Then, for each sex, we randomly draw 1000 times the number of bins in the region of interest bins from the genome-wide bins (208 bins) and we calculated the difference between the mean coverage of this random sample of bins and the region of interest. None of the comparison yielded a difference of 1 read or more (figure II.1C). In conclusion, there is no artefact due to coverage on the region of interest.

<u>X chromosome region chrX:47753028-47938680</u>
We extracted all X-linked position with a filter == PASS, outside the pseudo-autosomal regions.
The same analyses as above was perform. The mean coverage over 1000bp bins for the region of interest is presented on figure II.2A For males, the mean coverage is 7.50 (7.63 genome-wide, figure II.2B) and for female it is 14.55 (14.71 genome-wide, figure II.2B).
Then, for each sex, we randomly draw 1000 times the number of bins in the region of interest bins from the genome-wide bins (171 bins), and we calculated the difference between the mean coverage of this random sample of bins and the region of interest. None of the comparison yielded a difference of 1 read or more (figure II.2C). In conclusion, there is no artefact due to coverage on the region of interest.

**Figure II.1- Coverage analyses for the chromosome 17 region. A-** Mean coverage for males (red) and females (blue), the ribbon corresponds to the mean +- sd, for the region of interest. **B-** Distribution of the mean coverage for females and males for the whole chromosome 17 (top panels) and for the region of interest for females and males (bottom panel). **C-** Distribution of the difference between the mean coverage of the region of interest and the mean coverage of a randomly drawn set of 208 bins in the whole chromosome 17, for 1000 random samplings.

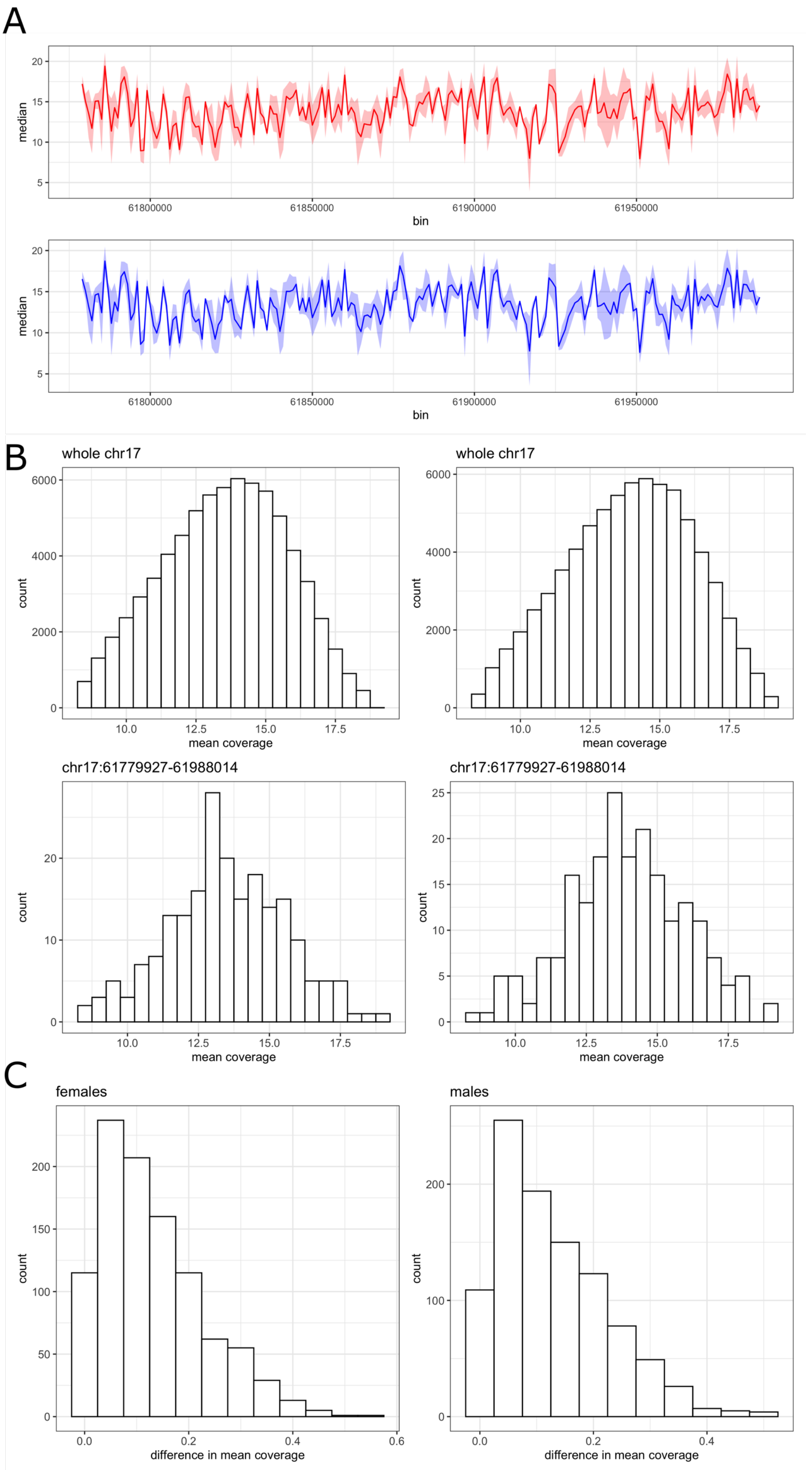

**Figure II.2- Coverage analyses for the X chromosome region. A-** Mean coverage for males (red) and females (blue), the ribbon corresponds to the mean +- sd, for the region of interest. **B-** Distribution of the mean coverage for females and males for the whole X chromosome without the PARs (top panels) and for the region of interest for females and males (bottom panel). **C-** Distribution of the difference between the mean coverage of the region of interest and the mean coverage of a randomly drawn set of 171 bins in the whole X chromosome, for 1000 random samplings.

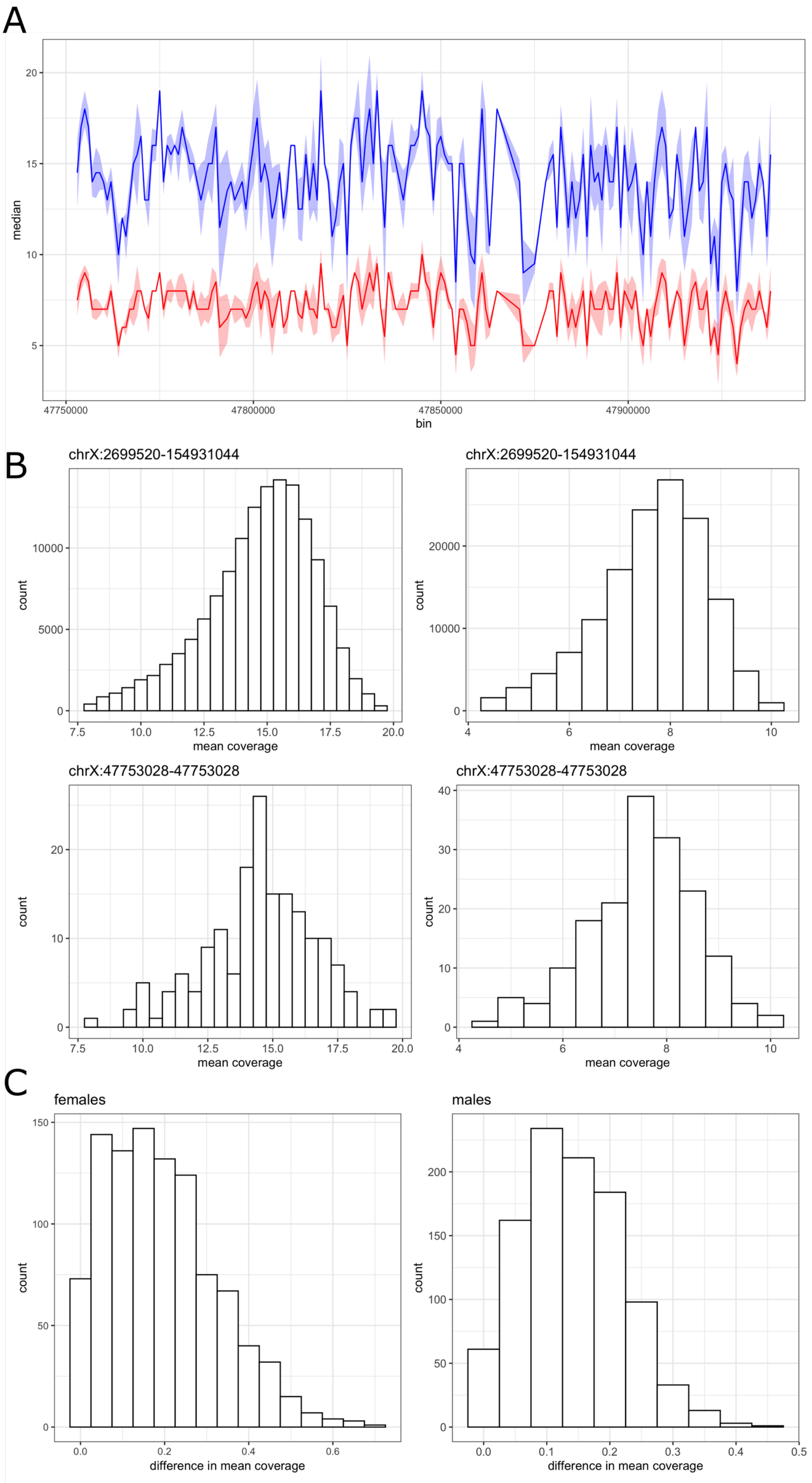

### III- Replication in the 1000Genomes populations

We extracted the phased genotypes of both candidate regions for all individuals included in the 1000 Genomes Phase 3. Then, we computed the matrix of distances between haplotypes within each populations, and performed a Multi Dimensional Scaling (MDS). We report here the results for European populations CEU (184 Utah resident with Northern and Western European Ancestry), TSI (112 Toscani in Italy), IBS (162 Iberian populations in Spain), GBR (107 British in England and Scotland) and FIN (105 Finnish in Finland).

For the chromosome 17 region, the three haplotypes are present in all European populations in the 1000Genomes dataset (Figure III.1). For the X-linked region, we can see individuals bearing at least 3 out of the 4 haplotypes detected in GoNL (haplotype 1, 2 and 4, Figure III.2).

**Figure III.1**- Multi Dimensional Scaling of the region of chr17:61779927-61988014 for the European population of 1000 Genomes. CEU (184 Utah resident with Northern and Western European Ancestry), TSI (112 Toscani in Italy), IBS (162 Iberian populations in Spain), GBR (107 British in England and Scotland) and FIN (105 Finnish in Finland).

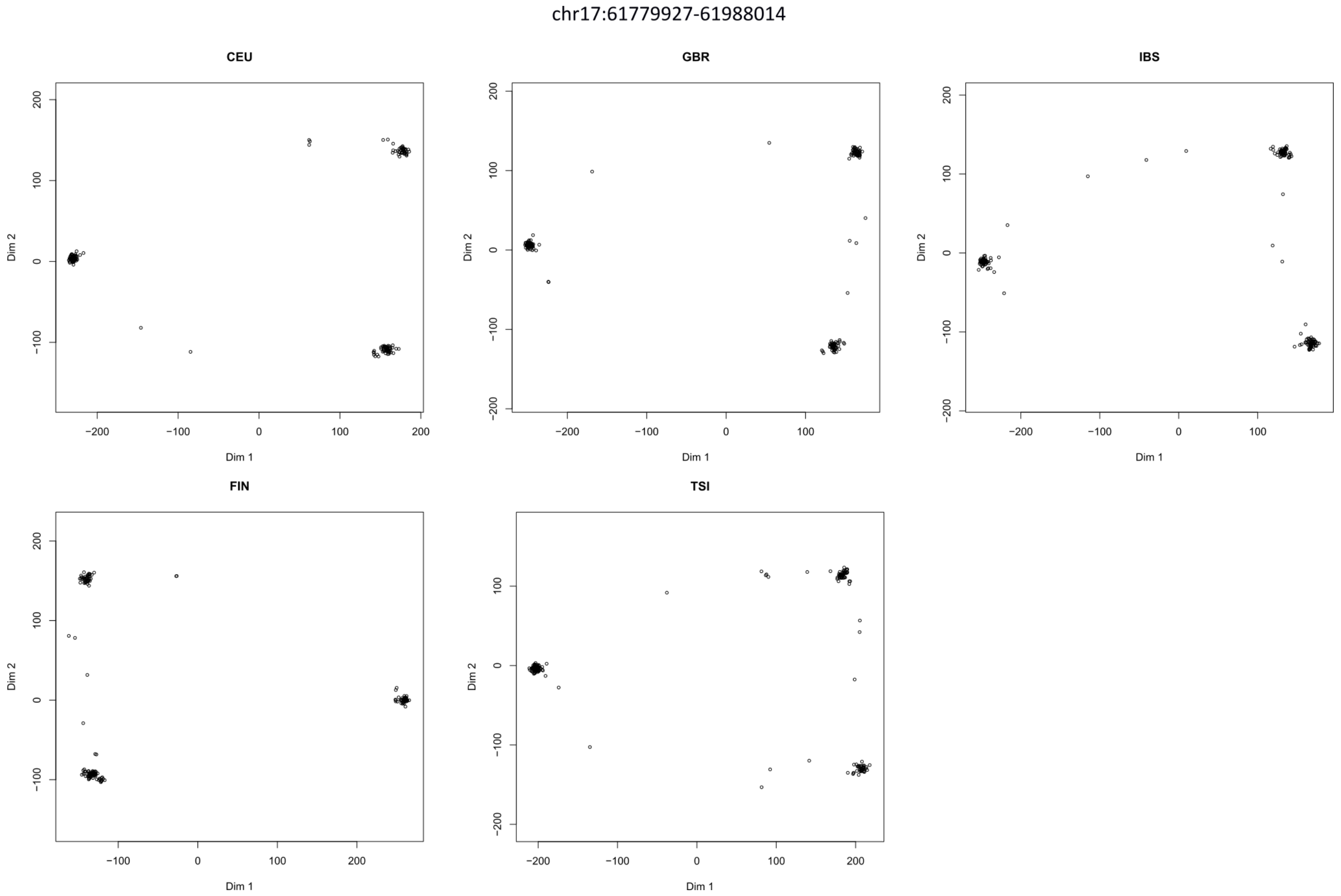

**Figure III.2**- Multi Dimensional Scaling of the region of chrX:47753028-47938680 for the European population of 1000 Genomes. CEU (184 Utah resident with Northern and Western European Ancestry), TSI (112 Toscani in Italy), IBS (162 Iberian populations in Spain), GBR (107 British in England and Scotland) and FIN (105 Finnish in Finland).

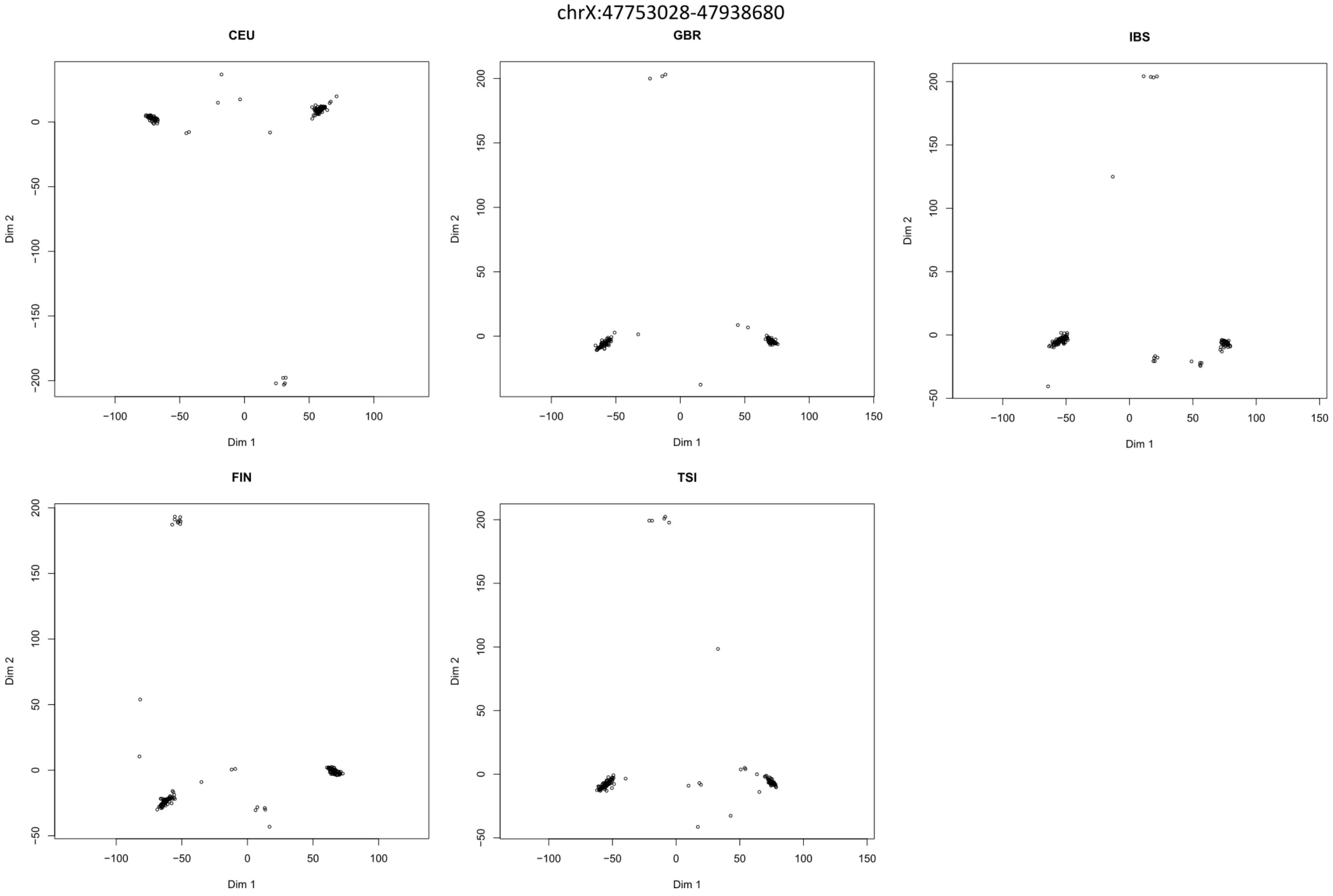

### SUPPLEMENTARY FIGURES

**Figure S1- Genome-wide analyses A-** Genomic localization of the Sex Antagonistic (red), Sex Limited (blue), Sex Differential (green) and Mixed (grey) transmission distortion regions.

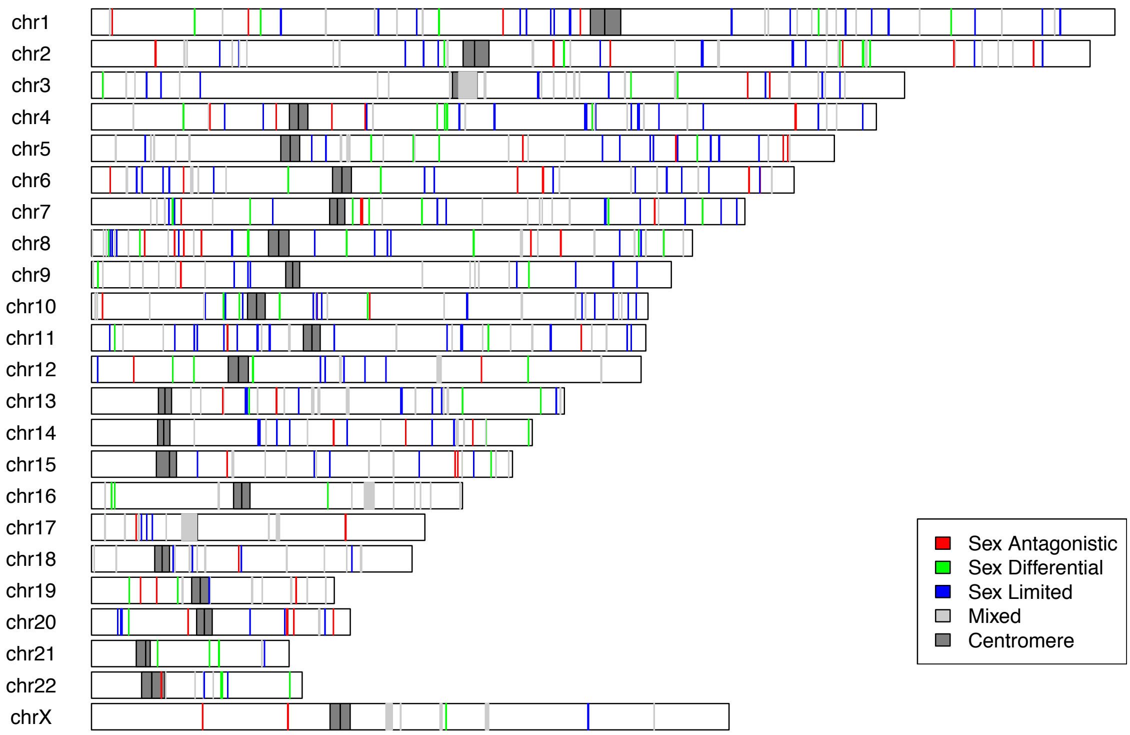


**Figure S2: Histogram of the thinned p-values** for 5272 SNPs that are 500kb apart.

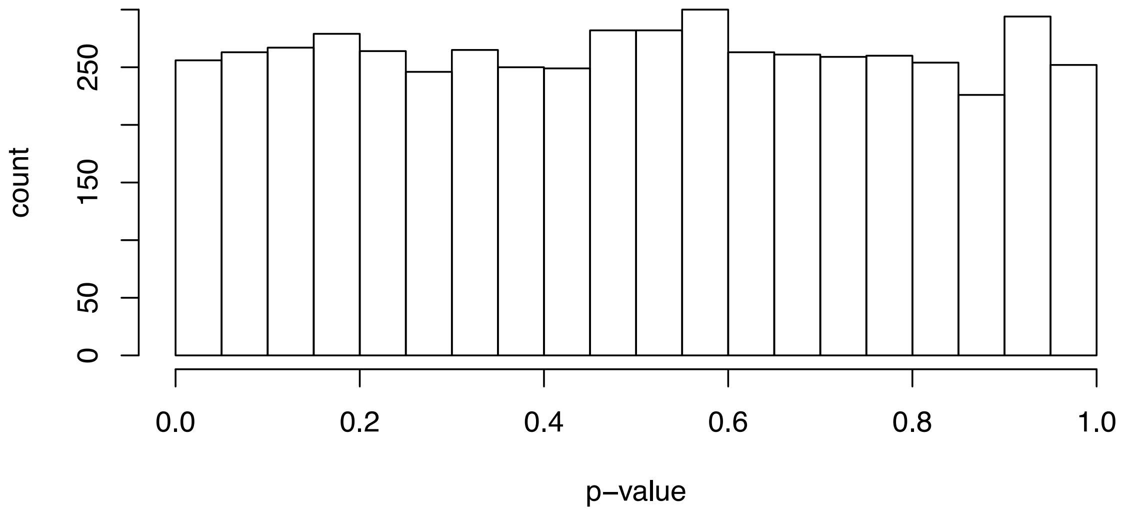

**Figure S3 –** Intersexual $F_{ST}$ distributions in both parents and offsprings in 248 trios for the three types of TD regions (in red) and for random regions with matching heterozygosity, length and number of SNPs (black).

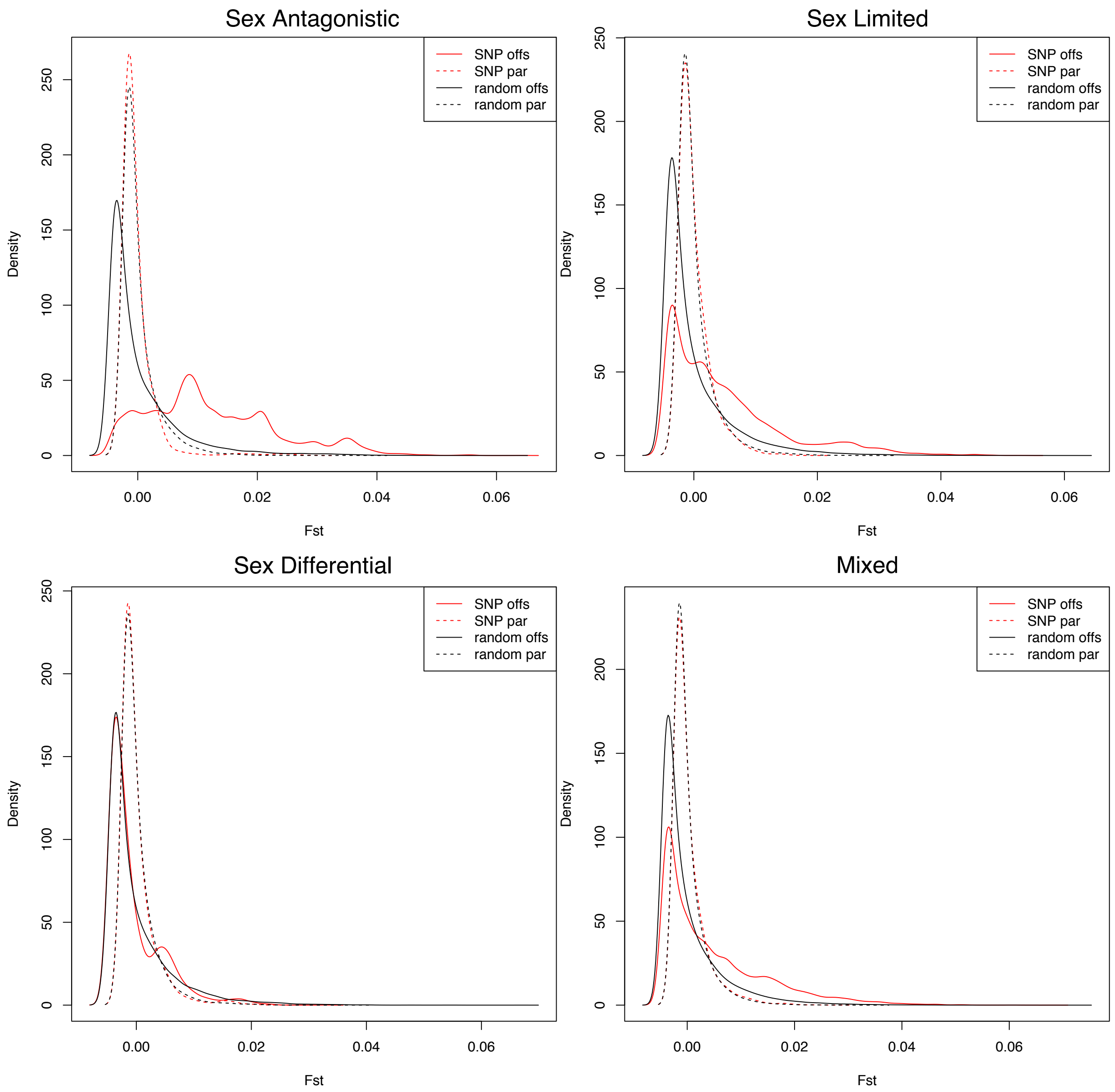

**Figure S4**- Genetic divergence between the three haplotypes discovered on region chr17:61779927-61988014 (208 kb): each line represents an individual and each column represent a SNP. A SNP is colored in grey if its genotype is the reference allele and in blue if it is the alternative allele.

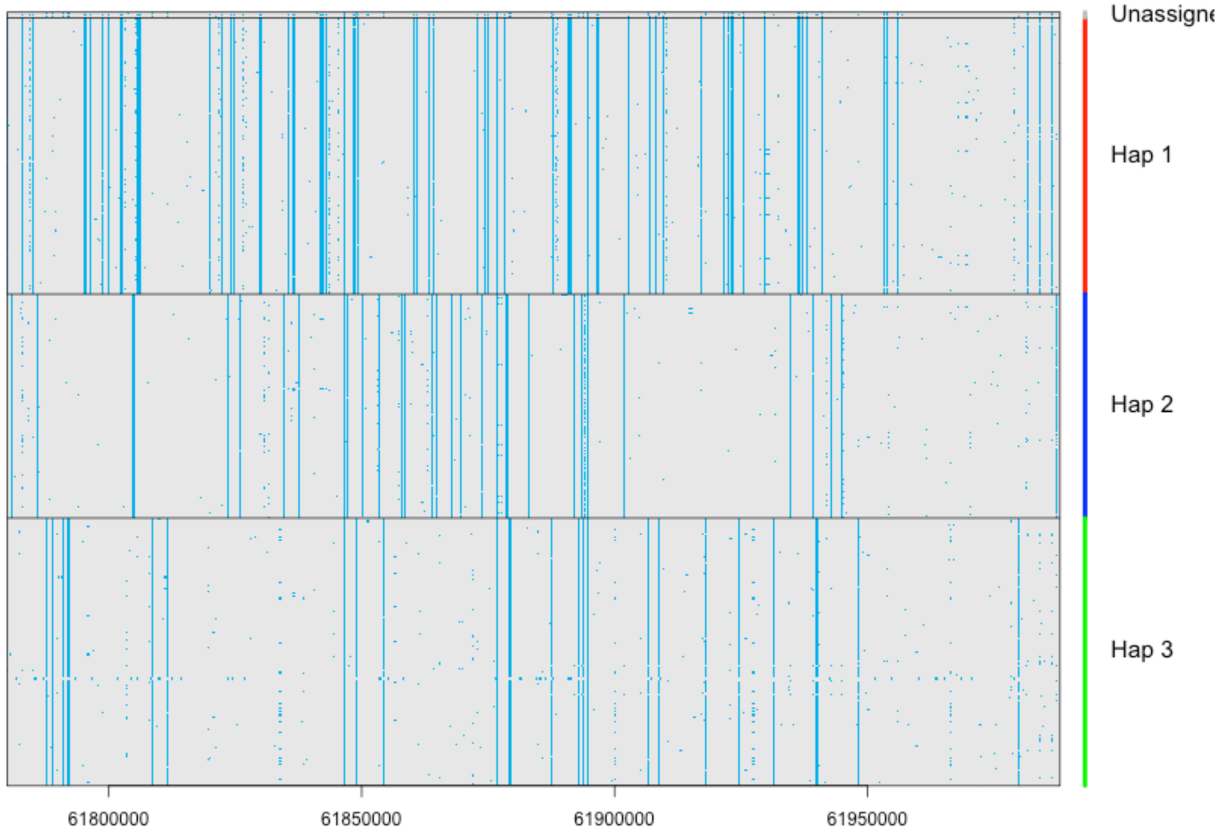

**Figure S5**- Genetic divergence between the four haplotypes discovered on region chrX:47753028-47938680 (186 kb): each line represents an individual and each column represent a SNP. A SNP is colored in grey if its genotype is the reference allele and in blue if it is the alternative allele.

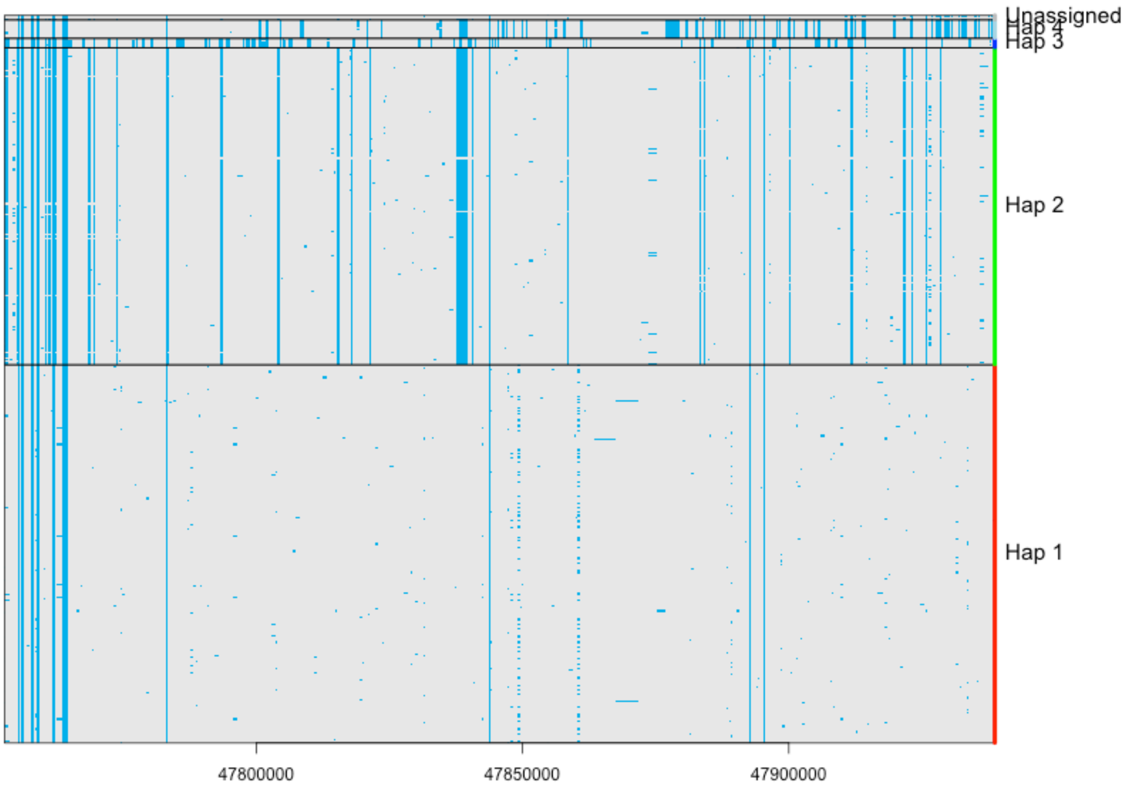

**Figure S6- MDS of the genetic distances between all individuals** (first two dimensions) for autosomes (1 million SNP randomly drawn, first draw) and the X chromosome (first repetition). The blue squares are females and the red circles are males.

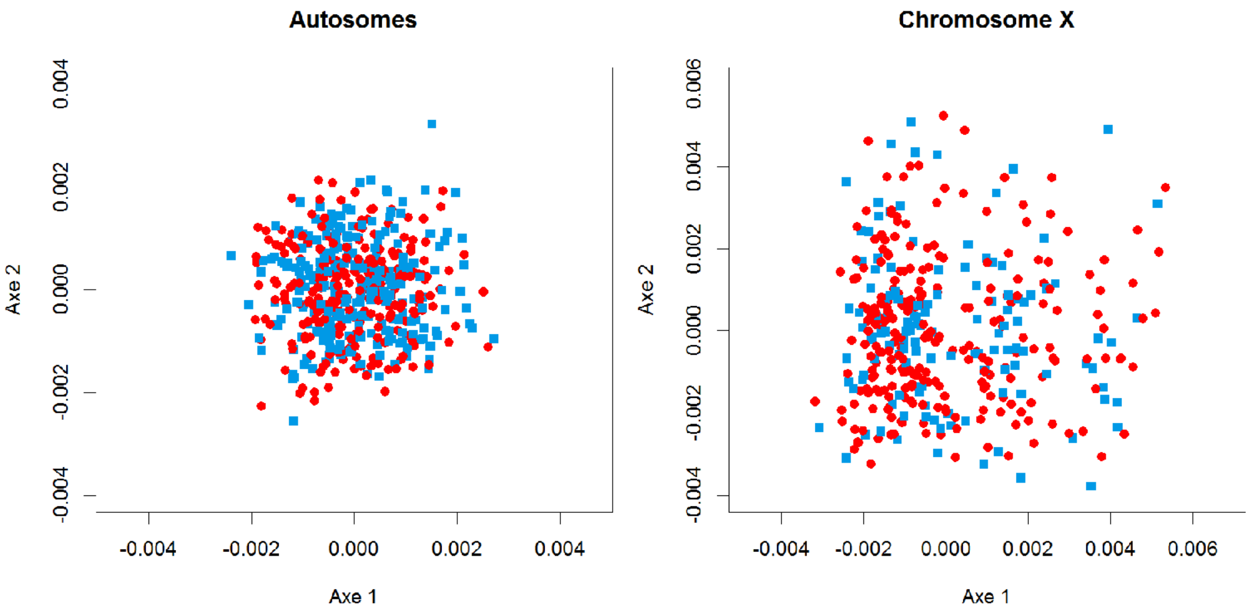

**Figure S7- Distribution of the age at sampling** for mothers (median=61), fathers (median=63), daughters (median=35) and sons (median=34).

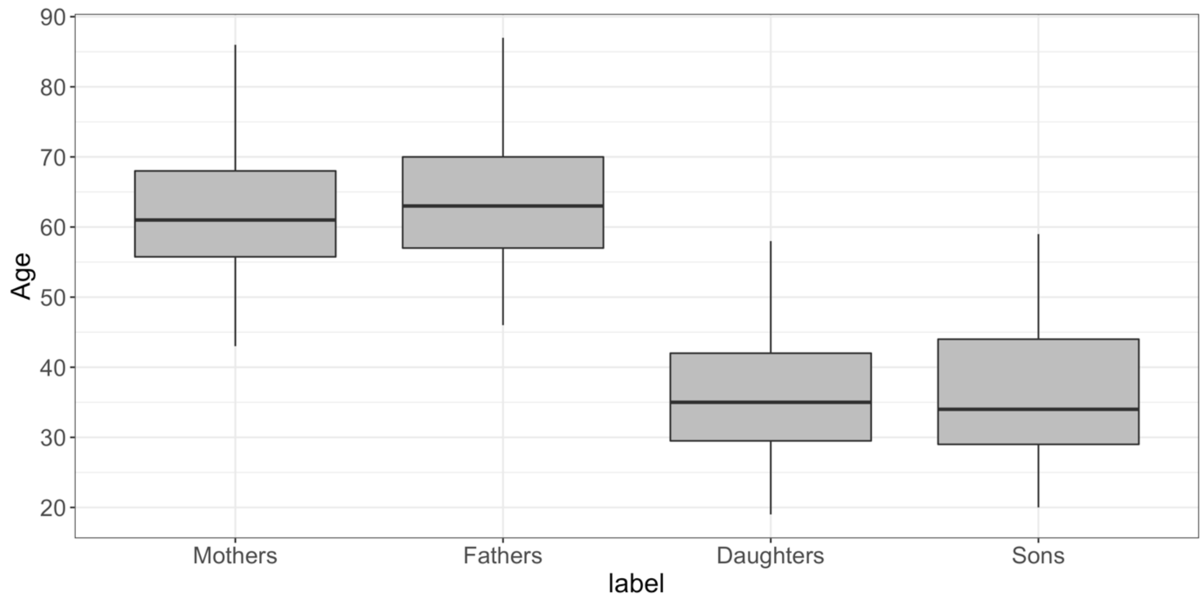

**SUPPLEMENTARY TABLES**

**Table S2 -** Summary of the enrichment found for the list of genes present in SD TD regions. For GO Biological Process and Human Gene Atlas, the enrichment is based on biological functions. For both GTEX and Jensen Tissue databases, the enrichment is based on which tissues the genes are expressed in.

| Database | Term | Overlap | P-value | Adjusted P-value | Z-score | Combined Score | Genes |
|---|---|---|---|---|---|---|---|
| **GO Biological Process** | choline catabolic process (GO:0042426) | 2/7 | 2.88E-04 | 5.14E-02 | -2.81 | 22.91 | DMGDH;BHMT |
| | L-methionine salvage (GO:0071267) | 2/8 | 3.83E-04 | 5.14E-02 | -2.63 | 20.66 | BHMT;BHMT2 |
| | cellular biogenic amine catabolic process (GO:0042402) | 2/9 | 4.91E-04 | 5.14E-02 | -2.87 | 21.88 | DMGDH;BHMT |
| **Human Gene Atlas** | Optic nerve hypoplasia (HP:0000609) | 2/24 | 3.63E-03 | 2.22E-01 | -2.32 | 13.02 | DDHD2;FGFR1 |
| **GTEX Tissue Expression Profile down** | GTEX-VJYA-0126-SM-4KL1P_spleen_male_60-69_years | 6/304 | 9.81E-04 | 1.00E+00 | -1.88 | 12.99 | DMGDH;BHMT;BHMT2;NDNF;PPL;FGFR1 |
| | GTEX-PW2O-0126-SM-48TC8_spleen_male_20-29_years | 6/311 | 1.10E-03 | 1.00E+00 | -1.82 | 12.39 | DMGDH;BHMT;BHMT2;NDNF;PTCHD4;PPL |
| | GTEX-POMQ-0126-SM-48TD6_spleen_female_20-29_years | 5/341 | 9.12E-03 | 1.00E+00 | -1.73 | 8.12 | DMGDH;BHMT2;NDNF;PTCHD4;PPL |
| | GTEX-UPK5-1626-SM-4JBHI_spleen_male_40-49_years | 5/353 | 1.05E-02 | 1.00E+00 | -1.77 | 8.04 | DMGDH;BHMT2;NDNF;BCAT1;PPL |
| **GTEX Tissue Expression Profile up** | GTEX-T5JW-1426-SM-4DM5Q_esophagus_female_20-29_years | 7/621 | 8.62E-03 | 1.00E+00 | -1.68 | 7.97 | BHMT;BHMT2;LSM1;CCDC102A;LETM2;GLYR1;FGFR1 |

**Table S3-** Summary of the enrichment found for the list of genes present in SL TD regions. For GO Biological Process and Human Gene Atlas, the enrichment is based on biological functions. For both GTEX and Jensen Tissue databases, the enrichment is based on which tissues the genes are expressed in.

| Database | Term | Overlap | P-value | Adjusted P-value | Z-score | Combined Score | Genes |
|---|---|---|---|---|---|---|---|
| **GO Biological Process** | tRNA catabolic process (GO:0016078) | 2/7 | 1.52E-03 | 2.54E-01 | -3.11 | 20.19 | EXOSC9;EXOSC8 |
| **Human Gene Atlas** | Goiter (HP:0000853) | 3/42 | 5.70E-03 | 3.92E-01 | -2.16 | 11.16 | TG;EYA1;SDHD |
| | Recurrent bacterial skin infections (HP:0005406) | 2/13 | 5.45E-03 | 3.92E-01 | -2.05 | 10.68 | NCF2;LYST |
| **GTEX Tissue Expression Profile down** | GTEX-S3XE-0626-SM-4AD6B_spleen_male_50-59_years | 8/308 | 5.47E-03 | 1.00E+00 | -1.89 | 9.86 | GRIA1;ALDH1L1-AS2;EYA1;ELOVL2;LAMA4;IPP;KCNK2;ULBP3 |
| **GTEX Tissue Expression Profile up** | GTEX-POYW-1226-SM-2XCEP_lung_male_60-69_years | 27/2090 | 2.21E-02 | 1.00E+00 | -1.68 | 6.41 | GRIA1;USP53;SYCP2L;NR3C1;LYST;ORC4;HERC3;CHORDC1;EPC1;GPBP1L1;SUPT20H;HLA-V;UBQLN4;TRPC3;API5;TPK1;IRAK3;ARPC5;FAM13A-AS1;TTC17;WDR11;ADAT2;PHF20L1;RIT1;ARHGEF2;RBMS2;CSNK1A1L |

**Table S4 –** Summary of the enrichment found for the list of genes present in mixed TD regions. For GO Biological Process and Human Gene Atlas, the enrichment is based on biological functions. For both GTEX and Jensen Tissue databases, the enrichment is based on which tissues the genes are expressed in.

| Database | Term | Overlap | P-value | Adjusted P-value | Z-score | Combined Score | Genes |
|---|---|---|---|---|---|---|---|
| **GO Biological Process** | positive regulation of protein localization to cell periphery (GO:1904377) | 7/47 | 4.84E-05 | 7.99E-02 | -1.47 | 14.57 | DPP10;PRKCI;EPB41L2;RANGRF;STX4;EPHA3;PLS1 |
| **GTEX Tissue Expression Profile down** | GTEX-T5JW-0326-SM-4DM6J_fallopian tube_female_20-29_years | 7/117 | 1.11E-02 | 1.00E+00 | -2.03 | 9.16 | ACBD5;PPFIBP2;STX19;DIRAS2;DEPDC1;NEB;SLC12A6 |
| **Mammalian Phenotype** | MP:0011096_embryonic_lethality_between_implantation_and_somite_formation,_complete_penetrance | 12/250 | 6.23E-03 | 6.62E-01 | -3.64 | 18.51 | CDH1;CHEK1;TOP3A;SETD1A;NMT1;KAT8;STX4;PFAS;MET;KPNB1;THAP11;ATR |

**Table S5-** Transmission counts of the three haplotypes detected in the region chr17:61779927-61988014, for heterozygous parents only. For example, haplotype one (186 cases) was transmitted 51 times out of 76 times to sons, and 44 times out of 110 times to daughters (significance assessed with exact Fisher tests).

| **Parents** | | **Transmission** | | | |
|---|---|---|---|---|---|
| haplotype | counts | to sons | pvalue | to daughters | pvalue |
| 1 | 186 | 51 / 76 | $\mathbf{3.84x10^{-3}}$ | 44 / 110 | $\mathbf{4.48x10^{-2}}$ |
| 2 | 164 | 21 / 54 | $1.34x10^{-1}$ | 58 / 110 | $6.34x10^{-1}$ |
| 3 | 184 | 27 / 68 | $1.14x10^{-1}$ | 66 / 116 | $1.63x10^{-1}$ |

**Table S6-** Distance between the three haplotypes detected in the region chr17:61779927-61988014 in terms of mean number of SNPs. In parenthesis is the mean percentage distance between all individuals (total number of SNPs is 1291).

| **distance** | **1** | **2** | **3** |
|---|---|---|---|
| **1** | 0 | | |
| **2** | 167.66 (13.00%) | 0 | |
| **3** | 151.78 (11.76%) | 84.59 (6.55%) | 0 |

**Table S7-** Transmission counts of the four haplotypes detected in the region chrX:47753028-47938680, for heterozygous parents only. For example, haplotype one (114 cases) was transmitted 18 times out of 45 times to sons, and 44 times out of 69 times to daughters (significance assessed with exact Fisher tests).

| **Parents** | | **Transmission** | | | |
|---|---|---|---|---|---|
| haplotype | counts | to sons | pvalue | to daughters | pvalue |
| 1 | 114 | 18 / 45 | 2.33E−01 | 44 / 69 | **2.95E−02** |
| 2 | 111 | 26 /43 | 2.22E−01 | 24 / 68 | **2.05E−02** |
| 3 | 7 | 1 / 2 | 1 | 2 / 3 | 1 |
| 4 | 8 | 2 / 4 | 1 | 3 / 4 | 6.25E−01 |

**Table S8-** Distance between the four haplotypes detected in the region chrX:47753028-47938680 in terms of mean number of SNPs. In parenthesis is the mean percentage distance between all individuals (total number of SNPs is 575).

| **distance** | **1** | **2** | **3** | **4** |
|---|---|---|---|---|
| **1** | 0 | | | |
| **2** | 40.61 (7.06%) | 0 | | |
| **3** | 44.13 (7.67%) | 41.4 (7.20%) | 0 | |
| **4** | 103.77 (18.04%) | 118.93 (20.68 %) | 146.04 (25.40 %) | 0 |
